# Language deficits in schizophrenia and autism as related oscillatory connectomopathies: an evolutionary account

**DOI:** 10.1101/044198

**Authors:** Elliot Murphy, Antonio Benítez-Burraco

## Abstract

Schizophrenia (SZ) and autism spectrum disorders (ASD) are characterised by marked language deficits, but it is not clear how these arise from gene mutations associated with the disorders. Our goal is to narrow the gap between SZ and ASD and, ultimately, give support to the view that they represent abnormal (but related) ontogenetic itineraries for the human faculty of language. We will focus on the distinctive oscillatory profiles of the SZ and ASD brains, in turn using these insights to refine our understanding of how the brain computes language by exploring a novel model of linguistic feature-set composition. We will argue that brain rhythms constitute the best route to interpreting language deficits in both conditions and mapping them to neural dysfunction and risk alleles of the genes. Importantly, candidate genes for SZ and ASD are overrepresented among the gene sets believed to be important for language evolution. This translational effort may help develop an understanding of the aetiology of SZ and ASD and their high prevalence among modern populations.

## 1 Introduction

Schizophrenia (SZ) and autism spectrum disorders (ASD) are pervasive neurodevelopmental conditions entailing several (and severe) social and cognitive deficits, including language problems (Bailey *et al*. 1996, Tager-Flusberg et al. 2005, van Os and Kapur 2009, Stephane et al. 2014). These deficits boil down to the impairment of basic cognitive functions and abnormal brain developmental and wiring paths during growth, which result in their distinctive neurocognitive profiles (Li et al. 2009, Stefanatos and Baron, 2011, Bakhshi and Chance 2015, Cannon 2015). Ultimately, both conditions result from the complex interaction between genetic, epigenetic and environmental factors. Recent advances in genome-wide technology have provided a long list of candidate genes for SZ and ASD which point to specific pathways and neural mechanisms that are seemingly dysregulated in them (Karam et al. 2010, Bennet et al. 2011, Flint and Munafò 2014, Jeremy Willsey and State, 2014, Geschwind and State, 2015). Nonetheless, the gap between genes, the pathophysiology of SZ and ASD, and their characteristic cognitive and linguistic profiles still remains open.

The goal of the paper is to contribute to the bridging of the gap between the genome and the abnormal linguistic cognition seen in ASD and SZ. We believe that this goal can be more easily (and more reliably) achieved if both conditions are examined together, instead of focusing on one or the other separately. SZ and ASD have been hypothesised to be opposite poles of a continuum of modes of cognition also encompassing typically-developing (TD) cognition, and their opposed natures can be tracked from brain structure and function to neurodevelopmental paths, to cognitive abilities (see Crespi and Badcock 2008 for review). Although several theories have been posited to account for this opposition, we wish to explore a novel possibility that builds on two different, but still related aspects: how the brain processes language, particularly through the perspective of brain rhythms, and how this human-specific ability evolved. We expect this novel approach will help achieve a better understanding of the aetiology of these two conditions, but also of language processing in the brain and of language evolution in the species.

Both SZ and ASD are characterized by abnormal patterns of neural oscillations (Moran and Hong 2011, Tierney et al., 2012, Pittman-Polleta et al. 2015), also during language processing (e.g. Braeutigam et al. 2008, Angrilli et al. 2009, Xu et al. 2012, Buard et al. 2013, Xu et al. 2013, Jochaut et al. 2015). Because brain rhythms are heritable components of brain function (Linkenkaer-Hansen et al. 2007, Hall et al, 2011) and because they are connected to some computational primitives of language (see Murphy 2015a for discussion), we believe that anomalous profiles of brain rhythmicity can help explain (and not just describe) language deficits in both diseases and map them to neural dysfunction and risk alleles of the genome (see Benítez-Burraco and Murphy 2016 and Murphy and Benítez-Burraco 2016 for an attempt). Interestingly, these profiles have proven to be disorder-specific (Buzsáki and Watson 2012), to the extent that cognitive disorders can be regarded as abnormal instances of the species-specific normal profile of brain rhythmicity, that is, as *oscillopathic* conditions. Interestingly too, because the hierarchy of brain rhythms has remained significantly preserved during evolution (Buzsáki et al. 2013), the human pattern of brain activity can be conceived of as a slight variation of a universal syntax of brain oscillations, and particularly, as a slight variation of the rhythmic patterns observed in other primates and presumed in extinct hominins. The changes in the skull and the brain that occurred after our split from Neanderthals and Denisovans seemingly brought about a distinctive, human-specific mode of cognition, which enabled us to transcend (better than other species) the signature limits of core knowledge systems and thus go beyond modular boundaries (Mithen 1996, Spelke 2003, Carruthers 2006, Hauser 2009, Boeckx 2011, Wynn and Coolidge, 2011). As reasoned in Boeckx and Benítez-Burraco (2014a), our language-readiness (that is, our species-specific ability to learn and use languages) boils down to this enhanced cognitive ability, but also to its embedding inside the cognitive systems responsible for interpretation and externalization. Importantly, this involves the embedding of high frequency oscillations inside oscillations operating at a lower frequency (see Boeckx and Benítez-Burraco 2014a for details). Putting it differently, the emergence of our language-readiness resulted from the emergence of a new global neuronal workspace which entailed new patterns of long-distance connections among distributed neurons and, consequently, new patterns of brain rhythmicity (see also Dehaene et al. 1998 for details).

It has been noted that recently-evolved neural networks are more sensitive to damage because of their reduced resilience (Toro et al. 2010). Accordingly, we should expect that a novel, human-specific cognitive ability, and particularly, the distinctive neural mechanisms it relies on, including brain oscillation patterns, is preferably impaired in modern populations, resulting in aetiologically-related, high-prevalent cognitive conditions. Interestingly, besides showing quite opposite symptomatic profiles, as noted above, SZ and ASD show as well a high prevalence within human populations (Murray et al. 2012). Because we have a good knowledge of the genes affecting both conditions, but also of the genes that have changed after our split from extinct hominins (Pääbo, 2014, Zhou et al. 2015), it seems worth testing whether the same factors that prompted the transition from an ape-like cognition to our human-specific, language-ready cognition are involved in the aetiology of cognitive disorders encompassing language deficits, like SZ and ASD. Furthermore, because our mode of cognition may boil down to a species-specific oscillatory profile (Murphy 2015a), and because both SZ and ASD are characterised by abnormal rhythmic patterns, we expect that investigating the genes related to brain dysrhythmias in SZ and ASD and interpreting their distinctive language deficits as oscillopathic features will allow a better understanding of their aetiology (and will ultimately lead to the construction of successful endophenotypes that allow an earlier diagnosis and a better treatment of the affected populations). Likewise, this effort should contribute to a better understanding of the nature of the human faculty of language, set against an evolving dynamic model of mental computation, and of its origins, set against a reorganizational model of the evolution of cognition.

The paper is structured as follows. First, we provide a comparative account of language deficits in SZ and ASD and review the anomalies in brain structure attested in both conditions, particularly the dysfunctions observed during language processing, with a focus on brain rhythms. Afterwards, we advance a novel oscillopathic model of language that leads to an enhanced and neurobiologically robust understanding of linguistic computation in the brain and language deficits in SZ and ASD. The framework we develop extends the model in Murphy (2015a) to achieve greater descriptive scope over a number of linguistic phenomena and characteristics of SZ and ASD. We then progress to the genome. We first check which of the candidates for the evolution of language-readiness are related to SZ, to ASD, or to both, with a focus on genes known to be involved in brain rhythmicity. We will check if the same genes are involved in both conditions and if they show opposite expression profiles. Finally, we examine the oscillopathic nature of language deficits in SZ and ASD (and the oscillatory nature of language processing in the brain) from a broader, evolutionary-developmental (evo-devo) perspective. Accordingly, we will try to show that examining the evolution of human language benefits our understanding of the nature of linguistic problems in SZ and ASD and the prevalence of both conditions, and, conversely, that exploring the oscillopathic nature of language deficits in SZ and ASD will enhance our knowledge of the nature and evolution of language.

## 2 Language deficits in the oscillopathic brain: SZ and ASD face to face

As reviewed in Benítez-Burraco and Murphy (2016), individuals on the autism spectrum display a range of communicative and linguistic difficulties. Around one third of children with ASD display morphosyntactic problems (Tager-Flusberg and Joseph 2003), and individuals with ASD more broadly use fewer functional words than individuals with Downs syndrome (Tager-Flusberg et al. 1990). They also integrate semantic information differently from typically developing controls during the interpretation of syntactic structures (Eigisti et al. 2011), and they consolidate semantic knowledge from such structures in a distinct way. Other linguistic impairments include problems with binding, relative clauses, wh-questions, raising and passives (Perovic and Janke 2013). These deficits more generally speak to a deficit in procedural memory.

Turning to SZ, Murphy and Benítez-Burraco (2016) explored as well in detail how individuals with SZ exhibit a broad range of phonological, syntactic and pragmatics impairments which lead to the following general typology of deficits: problems of speech perception (auditory verbal hallucinations), abnormal speech production (formal thought disorder), and production of abnormal linguistic content (delusions) (see Stephane et al. 2014). These and other forms of language impairments in SZ have been claimed to arise from problems with semantic memory, working memory and executive functions (Kuperberg 2010). Most of the distinctive symptoms have in turn been claimed to emerge directly from language dysfunction (Hinzen and Rosselló 2015). Individuals with SZ also have difficulty externalising their thoughts, displaying discourse-level impairments (Andreasen et al. 1985), but they also crucially display syntactic impairments, including the use of fewer relative clauses, shorter utterances, and less clausal embedding (Thomas et al. 1987, Fraser et al. 1986). This final impairment also implies difficulties engaging in thoughts about other mental states, such as *John believes that Bill likes Mary* (Morice and McNicol 1986). Finally, schizophrenics have a range of problems with reference and propositional meaning; deictic and definite noun phrases have been shown to be impoverished compared to non-definites (McKenna and Oh 2005). Both the ASD and SZ brain, then, display problems with clausal embedding, relative clauses and binding, although individuals with ASD have a number of additional syntactic limitations. At the level of the computational system, the linguistic difficulties seen in ASD and SZ amount to problems with agreement and word movement.

Unlike the normal left-lateralization of language-related brain activity in fronto-temporal regions, individuals with SZ typically exhibit bilateralism and right-lateralization (Diederen et al. 2010). The ASD brain also displays a number of structural and functional differences relatives to TD individuals. But since Hahamy et al. (2015) correctly point to the wide idiosyncratic variation in these differences, and since Murphy (2015a) explored the potential for neural oscillations to provide a more fruitful way of establishing brain-language linking hypotheses, Benítez-Burraco and Murphy (2016) and Murphy and Benítez-Burraco (2016) opted to turn instead to brain dynamics as a way of explaining linguistic deficits in ASD and SZ.

Unlike the nature – though not necessarily the localization of – linguistic representations in the brain, we feel that linguistic operations are presently a strong candidate for neurolinguistic investigation. Following Murphy (2015a), Benítez-Burraco and Murphy (2016) and Murphy and Benítez-Burraco (2016), we will assume that set-formation (the taking of two objects, X and Y, and forming the set [X, Y]) is implemented via the *α* rhythm embedding a series of cross-cortical *γ* rhythms. The precise algorithmic nature of this process will be discussed below. We will assume also that the ‘chunking’ of these sets emerges through embedding these *γ* rhythms within a medial temporal lobe – specifically, a hippocampal – *θ* band. The linguistic operation of labeling (maintaining a set of items in memory before coupling it with another rhythm to form a unique derivational identity), under our model, requires the slowing down of *γ* to *β* followed by *β*-*α* coupling via a basal ganglia-thalamic-cortical loop. We will additionally maintain that this is the only human-specific operation of language (Murphy 2015b). Finally, we will assume that agreement operations (establishing featural covariance in *John seems to like the film* and *John seemed to like the films)* are implemented via cross-cortical evoked *γ*, with this rhythm being involved in attention and perceptual ‘feature binding’ (Sohal et al. 2009). Importantly, although visual and auditory input would lead to distinct processing mechanisms, these computational operations would be predicted to be abstracted away from specific modalities, and hence the present oscillatory model would apply irrespective of sensory input. These operations will form part of what is termed the ‘oscillome’ in Murphy (2016a), or the set of oscillatory mechanisms which give rise to particular operations in computational cognitive science.

The brain dynamics of linguistic computation in ASD display a number of similarities with that of the SZ brain (see Table 1), as predicted in the present review of their linguistic deficits, with both ASD and SZ displaying problems with embedding, agreement and movement. They also display a series of opposing rhythmic properties (see Table 1), which we will claim explains recent hypotheses that ASD and SZ amount to opposite disorders (see, for instance, Crespi and Badcock 2008).

**Table 1.**
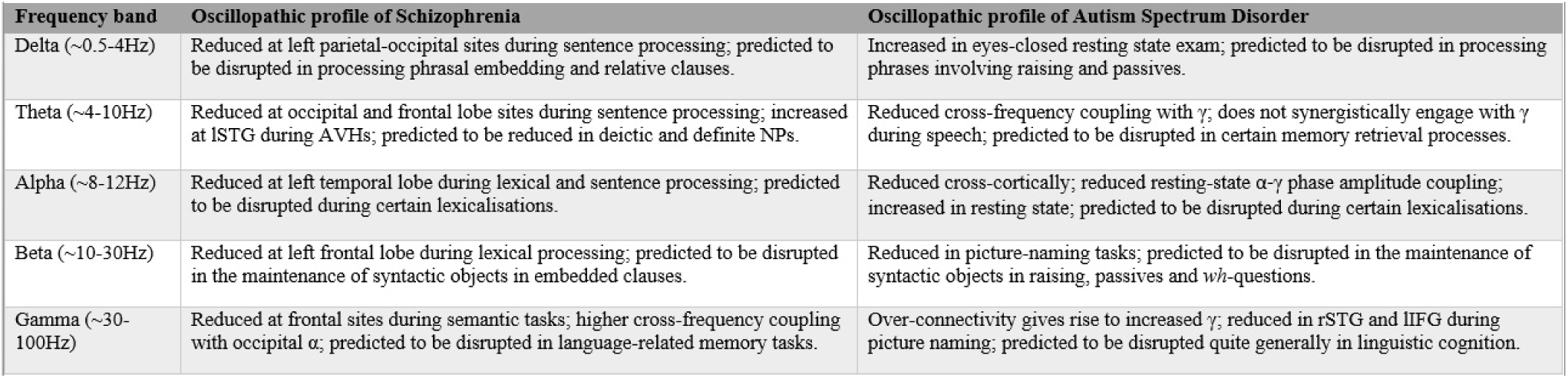
An oscillatory comparison between language deficits in SZ and ASD grouped by frequency range.

Both Kikuchi et al. (2013) and Rojas et al. (2008) reported significantly increased *γ* power for ASD individuals, with the former study additionally finding reduced cross-cortical *θ*, *α* and *β*. Xu et al.’s (2013) study of lexical decision in SZ also revealed reduced *α* and *β*, in the left temporal lobe and left frontal lobe respectively, suggesting problems with lexical selection and categorization. Reduced *α* and *β* was also documented in Moran and Hong’s (2011) and Uhlhaas et al.’s (2008) studies of SZ. Xu et al.’s (2012) sentence presentation task found reduced *α* and *β* in left temporal-parietal sites and reduced *θ* at occipital and right frontal lobe sites. Bangel et al. (2014) found reduced *β* during a number estimation task in ASD, while more general disruptions in rhythmic coordination have been frequently documented (see Benítez-Burraco and Murphy 2016 for a discussion of the possible connectomic reasons for these rhythmic disruptions, and Crossley et al. 2016 for a novel approach of ‘meta-connectomics’ which has the potential to comprehensively explore network connectivity in the SZ and ASD brain).

The ASD rhythmic profile also appears to frequently involve reduced *θ* during tasks requiring inter-regional synchronisation (Doesburg et al. 2013). Unusually long-lasting prefrontal and central *γ* has also been documented in ASD individuals during the interpretation of semantic incongruity (Braeutigam et al. 2008), possibly indicated the recruitment of a general search mechanism (*γ* rhythms) to replace the normal rhythmic processes (slow *γ*) responsible for retrieving and comparing semantic representations. Picture-naming tasks result in reduced *γ* and *β* in the left inferior frontal gyrus in ASD participants (Buard et al. 2013), while in SZ Uhlhaas and Singer (2010) point to reduced power in the same rhythms. Similarly for SZ, Ghorashi and Spencer’s (2015) visual oddball task led to reduced *β* at frontal, parietal and occipital sites. Resting-state *α*-*γ* coupling has also been found to be abnormal in children with ASD (Berman et al. 2015), while *θ*-*γ* coupling is severely impaired during speech processing in ASD (Jochaut et al. 2015). Jochaut et al. also found altered oscillatory connectivity between auditory and language cortices.

These close similarities in the rhythms putatively responsible for various linguistic deficits not only strengthen the current proposal that ASD and SZ should be characterized as oscillopathies, but they also support the present model of rhythmic language comprehension, proposed in Murphy (2015a). The social, pragmatic and resting-state oscillopathic literature seems to diverge substantially, however (see Hoptman et al. 2010, Spencer et al. 2004, Murias et al. 2007 and Uhlhaas and Singer 2010). Since the language-related deficits (in agreement, clausal embedding and so on) seem to map on to similar frequency bands and regions in ASD and SZ, it follows almost directly from this that the non-linguistic oppositions between the two disorders (as discussed in Crespi and Badcock 2008) must arise from a sub-group of these divergent connectomopathies. The next section will attempt to empirically demonstrate that particular linguistic computations appear to have the same oscillatory source across cognitive phenotypes, thereby lending support to a particular model of the human oscillome.

## 3 An SZ/ASD-driven oscillatory account of language processing

It was predicted in Benítez-Burraco and Murphy (2016) that due to the apparently preserved nature of the core linguistic computational operations in ASD and SZ, rhythmic variations in these disorders are likely to be the implementational source of representational and interface disruptions; that is, problems with information synchronisation between the rhythms and regions involved in phrase structure building and the external memory, conceptual and sensorimotor systems they are attempting to transfer structures to. We will here develop this idea in greater clarity. The SZ/ASD-informed oscillatory account of linguistic computation we will presently draw up should be seen as an implementation of the ‘Communication through Coherence’ hypothesis (Fries 2005, 2015), according to which rhythmic synchronization permits information transfer between neuronal groups. Even more broadly, we will follow Uhlhaas and Singer (2015) in interpreting the computational mind as arising out of a self-organizing neural system emerging from high-dimensional, non-linear dynamics.

Together with recent work into the oscillatory mechanisms responsible for working memory (such as Lisman and Jensen 2013), recent research such as Popov and Popova’s (2015) study of general cognitive performance in SZ may point towards a feasible model of linguistic computation more robust and fine-grained than the one proposed in Murphy (2015a) and discussed above. Popov and Popova revealed that individuals with SZ displayed higher *γ-α* cross-frequency coupling relative to typically developing controls, correlating with decreased working memory capacity. Higher coupling may result in smaller ‘gamma pockets’ of items retrievable from memory, with the disrupted coupling operations also impairing the normal retrieval order of these items, as discussed in Murphy and Benítez-Burraco (2016). We will hypothesize that this general coupling mechanism is employed in the service of phrase structure building, yielding the following components of linguistic computation:

i. The sequence of merged language-relevant features in a given derivational cycle at both lexical and phrasal levels.
ii. The capacity to search and compare agreeing features in both local and long-distance dependencies.

Contrary to Boeckx and Theofanopolou (2015), we see no way of exploring the oscillatory basis of linguistic computation without importing certain notions of what computations language seems capable of performing. Though we would ideally like to reach the point at which neurobiological investigations could impose direct constraints on linguistic theories, at the present state of the field we still find it necessary to consult the linguistic literature. For instance, if Narita (2014) is correct that phase-by-phase operations always proceed by initially merging an item from outside the derivation before applying any available movement, Agree and Transfer operations (with the latter set applying simultaneously), then this would provide a finer-grained rhythm from which to construct linking hypotheses between language and brain dynamics. Even if a phasal analysis is misguided, and the construct ‘phase’ is eliminated from the grammar – as proposed in suggestive work by Epstein, Kitahara & Seely (2014) – Narita’s ‘Merge + (Copy-formation + Transfer)’ constraint on structure building still ensures a level of systematic rhythmicity to the derivation. Recent work has also argued that linguistic structures can be labeled not only by standard categorial labels (NP, VP, etc.), but also by φ-features (Person, Number, Gender), as in [φ…α…[γ…β…]], expanding the implementational search range.

How would these proposals work in practice? We will assume that particular oscillatory coupling operations retrieve lexical features from widely distributed conceptual regions they are coupled with and transfer them to other systems oscillating at different frequencies. In order to understand how this takes place, we need to draw up a model of the interfaces. This amounts to particular gamma pockets of particular phasal properties being able to synchronise neural firing with other systems. Derivational feature-checking (e.g. φ-feature agreement followed by Q-feature agreement within the same cycle/phase) may arise from the particular sequence of items extracted within a given oscillatory cycle. Below is a hypothesized example of hippocampal *θ* extracting a series of features from distributed regions oscillating at low to middle *γ* – taking into consideration Lewis and Bastiaansen’s (2015) speculations about evoked high *γ* being implicated in the propagation of bottom-up prediction errors – with the numerical indices referring to the order of faster rhythms embedded within the slower rhythm and the ‘Q’ and ‘φ’ referring to the extracting featural items from memory (see Figure 1 for a schematic representation):

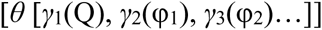

The series of feedforward *γ* rhythms employed in this model would be most prominently generated in supragranular cortical layers (L2/3) (Maier et al. 2010), while hippocampal *θ* would be produced by slow pulses of GABAergic inhibition as a result of input from the medial septum, part of a brainstem-diencephalo-septohippocampal *θ*-generating system (Vertes & Kocsis 1997). The crucial interactions between the hippocampus and medial prefrontal cortex (mPFC) necessary to focus attention on language-relevant features (considering the conclusions of Lara and Wallis 2015 on the role of prefrontal cortex in working memory, which stressed the centrality of attention rather than storage) may be mediated through an indirect pathway passing through midline thalamic nucleus reuniens (Jin and Maren 2016). The potential significance of the thalamus will be a theme returned to below, but we would like to suggest here that due to the aberrant functional coupling between the hippocampus and mPFC both during rest and working memory tasks in SZ (Lett et al. 2014) may result not only in the deficits in emotional regulation seen in SZ, but may also play a role in particular linguistic deficits involving the extraction of incorrect feature-sets (and, in conjunction, may yield problems with emotion-related language).

**Figure 1.**
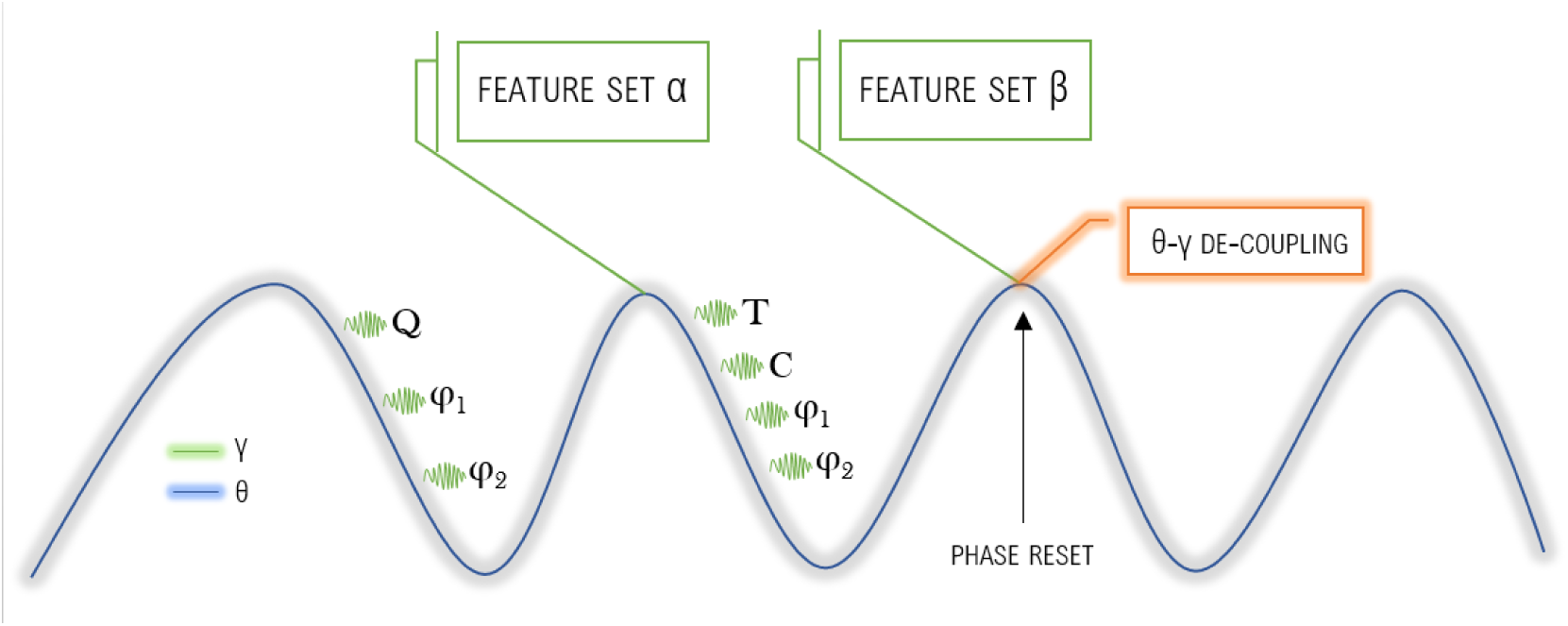
An idealised schema for the retrieval of a sequence of language-relevant features in a given derivational cycle, as proposed in Murphy (2016). ‘Q’ denotes Q-feature, ‘T’ denotes Tense feature, ‘C’ denotes Case feature, and ‘φ’ refers to φ-features (e.g. 1stP, Singular). After action potentials fire at the trough of each slow cycle and as inhibition reduces over the θ cycle, the most excited cluster/representation would be retrieved first via low-middle γ, followed sequentially by the second and third most excited. The principles of ordering and excitability which determine the composition of the feature-set (completed after the θ phase resets) remain to be worked out, but an approach similar to Jensen et al.’s (2012) model of visual attention may be fruitful, ultimately imposing neurobiological constraints on theories of linguistic feature-set construction. For instance, what determines the order of item extraction might be the connection strength between neuronal ensembles responsible for storing the representation in memory and/or their excitability. External constraints would also enter into determining the order of the temporal serialization of item extraction, with Ray and Maunsell (2015) noting that the coordination of γ phase across multiple, distant areas is difficult due to conduction delays, mediated by myelin thickness and nodal structure. A 5ms conduction delay, for instance, has been estimated to change the interactions of two coupled high γ oscillators from constructive to deconstructive interference, while delays less than 1m could alter the phase by 30° (Pajevic et al. 2014). Altered gene expressions effecting myelinization could consequently play a substantial role in altering oscillatory, and consequently linguistic, competence.

When a lexical item has a particular featural configuration such that is requires, for instance, a subject (as in Tense Phrases), this may be the result of a particular embedded rhythm requiring a particular set of subsequent cycles to activate the required representation. Under the present proposal, the online structure-building would crash and become uninterpretable only if this generic mechanism extracts the incorrect sequence of features. After each top-down, feedback *θ* cycle (in predictive coding terms; Friston 2005) extracts feature-sets from memory, the embedded fast *γ* would slow to the *β* band to be maintained in memory (with frontal *β* also likely playing a role in the regulation of cognitive control in the above hippocampal-mPFC network; Stoll et al. 2015). As the sentence is processed online, the cumulative *γ* items which decrease in power to *β* would consequently increase *β* power over regions responsible for storing and constructing linguistic representations such as BA44, BA45, BA47 and the left temporal lobe, incorporating the neural clusters charged by *γ* cycles into a system of online language comprehension. Each *γ*-matched cluster would therefore correspond to a node on the functional phrase structure network. Bottom-up *γ* would be able to rapidly adjust the ongoing set of lexical and phrasal representations via a familiar feedforward mechanism, updating the localised *θ* in the hippocampus and the more widely distributed inter-areal *β*, with the latter rhythm possibly being responsible for linking distinct cortical areas into what Bressler and Richter (2015) term NeuroCognitive Networks (NCNs); large-scale, self-organizing cortical networks. Bressler and Richter argue that *β* plays a dual role, being involved in NCN maintenance and the transferring of top-down signals to lower levels in the cortical hierarchy such as the *γ* range. We think that this perspective fits well with the standard requirement within linguistic theory that phrases must be labeled via two (for us, domain-general) sub-processes: object maintenance (holding the constructed set in memory) and property attribution (giving the set a syntactic identity independent of its components, e.g. forming a phrase from a noun and a verb yields a Verb Phrase), since *β* would be able to simultaneously maintain an object as a cognitive set (via its steady or increasing amplitude) and attribute a specific representational property to it (via top-down feedback and transferring prediction signals). Indeed, it is also possible that the distribution of syntactic categories in a sentence (C, T, *v*, D, and so on) is implemented through a similar procedure, albeit through far more distributed networks being responsible for maintaining the general featural properties of each category type; we would not expect a network of verbal representations to be smaller than the network responsible for storing one particular verb type, for instance.

Even though the connectomic and local network properties of the γ-oscillating regions responsible for storing featural representations would likely impose certain constraints over the order of item extraction within *β* and *θ* waves, given what has been outlined here we think it is also likely that a source of grammatical and feature-combination constraints arises from the top-down information transferred from these slower waves down to lower regions of the cortical hierarchy, such that the representations constructed by γ-itemized objects would pass through a general ‘filter’ at the θ-oscillating regions (imposing memory-related rules such as relevance and attentional priorities, in the spirit of Jensen et al.’s 2012 approach to the visual system’s prioritization of salient unattended stimuli via posterior α) and β-oscillating regions (imposing general cognitive set-constructing rules of efficiency) they are phase-locked to. This perspective is approximately in line with recent proposals that processing of syntactic complexity is distributed across the language system, rather than being narrowly localised purely within the left inferior frontal gyrus (Blank et al. 2016). A recent study of γ-band synchronization (GBS) in children also revealed greater GBS when a representation of two objects was held in memory during occlusion relative to one object, reflecting object maintenance demands rather than sensory processing and lending support to our hypothesis that *γ* power reflects the generation of item sequences from across cognitive domains (Leung et al. 2016). Medial temporal lobe theta rhythms increased when participants successfully link two distinct memories. These rhythms were synchronised with the mPFC, an area involved in storing knowledge networks. Increased hippocampal-mPFC *θ* coupling appears to be causally implicated in memory integration (Backus et al. 2016). The general cognitive processes used to resolve linguistic dependencies may also be used to resolve music and action sequences (Van de Cavey and Hartsuiker 2016), and future experimental work could address whether the oscillatory coupling mechanisms are also similar, perhaps as a result of the language system having ‘copied’ its generic embedding mechanism across to other domains via neural ensemble connections.

This rich interplay between distinct rhythms (which includes the role of *α* in synchronizing cross-cortical regions) would amount to a comprehensive ‘neural syntax’. The computational modeling in Kopell et al. (2011) suggests that *β* has the appropriate temporal characteristics to maintain information across typical short-term and working memory time-spans. Recent literature, reviewed below, lends support to this rhythmic and regional functional segregation. To illustrate, this theory permits a simple explanation for why a larger coherence in the 13-18 Hz range was found by Weiss et al. (2005) in response to syntactically complex object-relative clauses in comparison to less complex subject-relatives, since the former structures require a greater computational load in terms of feature-matching. Our perspective also allows the natural re-translation of the notion of a memory stack, with each slow rhythm operating as spatiotemporally distinct stacks of feature-sets retrieved by a sequence of faster rhythms.

Recent work suggests that a decree-se of left-temporal and left inferior frontal *α* reflects the engagement of higher-level language regions which pre-activate anticipated, predictable words. This anticipatory effect was also seen in the hippocampus (Wang et al. 2016). Other recent work points to the predictive role of *γ* alongside the role of *β* in maintaining existing cognitive states and objects in memory (Lewis et al. 2015), and these findings would be commensurable under an additional assumption that a generic phase-amplitude coupling mechanism embedding predicting/retrieving *γ* inside *α*. We believe that this may suggest that slow *α* is being used as a general prediction-retrieval mechanism which in turn synchronizes with hippocampal *θ* in the service of retrieving the successful set of featural representations which form the predicted word upon the completion of the *α* phase. When the brain jointly employs a number of these cross-frequency coupling operations in spatially overlapping networks (hippocampus, basal ganglia, thalamus, BA44, BA45, BA47) it may achieve a degree of unification between distinct hierarchical representations, e.g. syntactic and phonological representations would be unified into a coherent oscillatory structure, yielding core features of human language. The coupling of *β*, *γ*, *θ* and *α* at different time scales would allow the brain to flexibly achieve the rapid parsing of linguistic structures, with an average reader comprehending 250-300 words per minute (Rayner et al. 2012).

This theory of feature-set computation would explain why the higher *γ-α* cross-frequency coupling in SZ (resulting in smaller chunks of extracted representations) leads to an impoverished working memory capacity. We predict that future studies of the SZ oscillome focusing on language comprehension would deliver similar results, with the model in Figure 1 being compromised. Our approach (which amounts to a refined version of the model in Murphy 2015a) can also shed light on the findings of Bastiaansen et al. (2010). Their MEG study of licit sentences, sentences containing a syntactic violation, and randomised words revealed a gradual increase in low *β* (13-18 Hz) power during the first two conditions, which was disrupted at the point of syntactic violation (see Davidson and Indefrey 2007 for similar findings). In the present model, this would reflect the ongoing labeling (and re-labeling) of the structure, and we would predict that this slow *β* would be coupled with low and middle *γ* at the ensembles responsible for maintaining the feature-sets extracted for the purposes of constructing the online lexical and phrasal representations. This rhythm-based segregation of decomposed, primitive and generic linguistic sub-operations has the potential to explain a variety of necessary functions in language comprehension. Further evidence that *β* in particular is involved in featural agreement comes from Perez et al.’s (2012) study of ‘unagreement’ in Spanish, which showed that sentences containing an agreement violation but which nevertheless led to a grammatical sentence (in their case, a mismatch between the subject and verb’s Person features) led to decreased *β* power at the verb relative to other grammatical sentences which did not involve agreement violations. This has been interpreted in the literature as indicating simply that the language faculty is marking a change in cognitive set (Lewis et al. 2015), however we think that our oscillatory feature-set model permits a more computationally rigorous explanation. There is an additional possibility, emerging out of the work on *α* and visually induced *γ* by van Ede et al. (2015), that the rich combinatorial possibilities of feature-set composition and agreement relations seen in human language may also arise out of the diversity in oscillatory phase-relations between neighbouring regions, which are likely to be much greater in the larger and remarkably globular human braincase in comparison to the ‘spatial inequalities’ (Salami et al. 2003) between cortical and subcortical regions documented in less globular brains; a topic to be returned to below. As Ede et al. (2015: 1556) note, in humans ‘phase-relation diversity is a general property of neural oscillations that is restricted neither to the lower frequencies nor to periods outside of stimulation’. It is of particular interest that phase-relation diversity has been documented in low *θ* (Agarwal et al. 2014, van der Meij 2012) and *α* (Bahramisharif et al. 2013), possibly yielding the computationally flexible rate at which linguistic features can be merged and copied (Narita 2014). In conjunction with evolutionary concerns about globularization, other recent research suggests new directions for pursuing our hypothesis and establishing further brain-language linking hypotheses. Using Laplace Eingenmodes to analyse MRI and DTI data, Atasoy et al. (2016) showed that resting brain function is related to brain shape. The harmonic waves they observed appear to obey the same physical principles as other self-organising phenomena like tiger and zebra stripes or the patterns of vibrating sand; lending weight to Descartes’s original intuition that the brain is organised through principles of efficient causation, and not being incompatible with recent work in generative grammar which suggests that syntactic computation operates via principles of efficient computation (Narita 2014). One of the major goals of the current Dynamic Cognomics enterprise is consequently to build (and, ultimately, replace) the traditional syntactic trees of linguistic theory with ‘oscillomic trees’, following the general decompositionalist proposals in Murphy (2015c) (see Figure 2 for how this approach might relate to studies of SZ).

**Figure 2.**
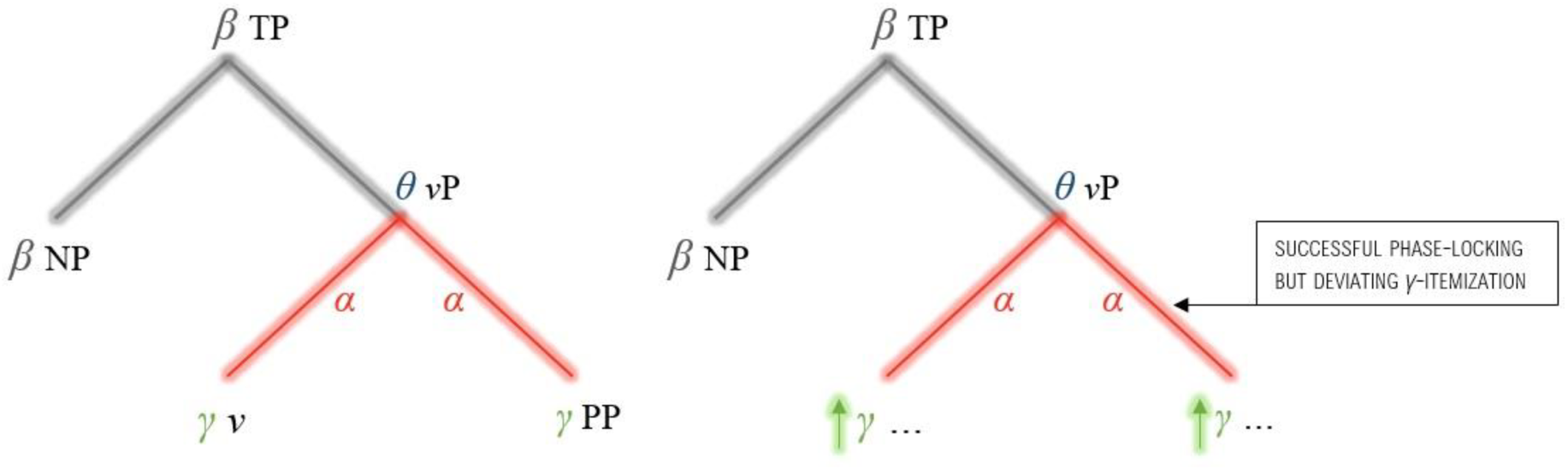
An oscillomic tree representing a typically developing oscillome (left) and an SZ one (right), in which the unusually fast γ in SZ during coupling with α leads to itemization sizes of an unsuitable frequency for successfully extracting particular linguistic representations (see Uhlhaas and Singer 2010, Popov and Popova 2015, Boeckx et al. 2016, Murphy and Benitez-Burraco 2016, Murphy 2016). ‘TP’ denotes Tense Phrase, ‘vP’ denotes Verb Phrase (e.g. ‘swam in the river’, ‘NP’ denotes Noun Phrase (e.g. ‘The man’, ‘John’), and ‘PP’ denotes Prepositional Phrase (e.g. ‘in the river’).

Related studies also point to an extension of the model in Figure 1 to account for local and long-distance filler-gap dependencies, which could be employed in experimental settings investigating the oscillome of the SZ and ASD brain when interpreting (or failing to interpret) agreement relations. Meyer et al. (2013) compared the oscillatory responses to local and long-distance dependencies between a subject and a verb in subject- and object-relative clauses in German. At the point of agreement resolution upon interpreting the verb, Meyer et al. discovered higher *β* for long-compared to short-distance dependencies. A possible explanation for this is that the higher working memory load implicated in long-distance dependencies necessitates the deployment of a higher frequency range in order to synchronize the activation of the ensembles implicated in the feature-sets of the filler and gap, with particular cellular clusters likely being pre-activated by the filler (since dependent elements in a sentence share a sub-set of their features). Since *γ* has been shown to be modulated by cloze probability (Wang et al. 2012), it may be that *β*-*γ* synchronization is the core (possibly the only) mechanism instrumental to the binding of feature-sets within phrase structures, with *γ* being responsible for semantic prediction and feature-binding (compositional meaning) and *β* responsible for syntactic feature-binding and object maintenance (labeling). This proposal becomes stronger when we acknowledge that the tissue these oscillations emerge from are partly based on the left inferior frontal gyrus, which for us is not the ‘seat of syntax’ as many have claimed, but rather an active memory buffer for the construction of lexical and phrasal feature-sets. Adopting these assumptions leads to a scenario in which oscillations may not only be the generic, domain-general mechanisms which allow networks of thousands of neurons to interact, but they may also function to orchestrate the construction of linguistic structures. It would also follow from the present hypothesis that a characteristic feature of human syntax, long-distance dependencies, arises from the novel setting of a generic oscillatory mechanism operating on an extensive range of brain regions as a result of a globularized braincase (see the following sections for a discussion of these evolutionary concerns).

In the same way that O’Keefe and colleagues have successfully localised the neural ensembles responsible for particular spatial representations in mice, future hemodynamic neuroimaging and magnetoencephalographic studies of human language should attempt to localise the ensembles responsible for particular featural representations (Tense, Person, Number, Gender, Nominal, Verbal, and so on). These will doubtless be comparatively more widely distributed than the cells responsible for vector formation in the mouse hippocampus, but the dynamic coupling capacities of the oscillome would hypothetically be employed to trigger the activation of such highly specific (and functionally specified) clusters. In addition, although we find the recent work of Poeppel and colleagues illuminating and promising, we also feel that this top-down experimental approach has a number of limitations. For instance, while Ding et al. (2016) delineated the rhythms responsible for ‘packaging’ particular linguistic constituents, the functional role of these rhythms in cognition more generally needs to be explored in much greater detail. This approach would ultimately require the establishment of novel linking hypotheses between the experimental work on the entrainment of particular rhythms to linguistic constituents and broader research into the oscillatory nature of working memory, attention, and consciousness.

Language should no longer be seen as a Fodorian module, since under this view it is rather a cross-modular system arising from the interfacing of diverse oscillatory mechanisms performing low-level, generic functions. Empirically testing this model will require, for instance, MEG and EEG scanning of individuals with SZ and ASD during specific language tasks, such as the comprehension of a range of clausal embeddings, long-distance agreement cases (both interpretable and illicit), local and long-distance word movement, and other grammatical constructions which could reveal the phase-relation and coupling properties of a linguistically impaired oscillome.

## 4 SZ- and ASD-related genes and the evolution of language-readiness

### 4.1. Common candidates for SZ and ASD?

Having established how the oscillome of individuals with SZ and ASD appears to function, we now turn to the genetic basis of these oscillopathic conditions. As noted in the introduction, the number of candidate genes for SZ and ASD has grown exponentially over the years. Many of these genes map on to specific pathways and brain functions that are associated with susceptibility to these conditions. For instance, knockdown of key candidates for ASD (*Mecp2*, *Mef2a*, *Mef2d*, *Fmr1*, *Nlgn1*, *Nlgn3*, *Pten*, and *Shank3*) in mouse primary neuron cultures alters the expression of genes mapping to pathways associated with neurogenesis, long-term potentiation, and synaptic activity (Lanz et al. 2013). Likewise, co-expression analyses of nearly 500 candidates for this condition across several brain regions spanning prenatal life through adulthood have uncovered modules of co-expressed genes that are related to synapse formation and elimination, protein turnover, and mitochondrial function (Mahfouz et al. 2015).

Similarly, analyses of GWA data related to SZ have revealed enrichment in several neural pathways and functions, like neurotrophin signaling pathway and synaptosome assembly (Chang et al. 2015), corticogenesis and neurotransmitter homeostasis (Papaleo et al. 2012), and aspects of postsynaptic density at glutamatergic synapses (Hall et al. 2015).

Importantly, some of the candidates for SZ and ASD seem to play a role in the maintenance of the adequate balance between neuronal inhibition and excitation and thus, in brain rhythmicity. Among them one finds genes that encode ion channels or protein associated with them, proteins involved in neurotransmitter activity, and proteins related to synaptogenesis. Hence, promising candidates for SZ are *CACNA1I*, which encodes a calcium channel; *CACNA1C*, encoding one of the subunits of the Cav1.2 voltage-dependent L-type calcium channel; *DPP10*, which encodes a membrane protein that binds specific potassium channels; and *CNTNAP2*, associated with K+ voltage-gated channels (all of which are reviewed in Murphy and Benítez-Burraco 2016). In turn, *Nav1.1*, which encodes a sodium channel (reviewed in Benítez-Burraco and Murphy 2016), is a candidate for ASD. Likewise, several genes encoding neurotransmitter receptors have been associated with abnormal oscillation patterns in schizophrenics and autistics, and ultimately to the language problems they exhibit (see Figures 1 and 2). Specifically, among the candidates for SZ one finds genes encoding receptors for serotonin, like *HTR1A*, and NMDA, like *GRIN2A*, which causes epilepsy-aphasia spectrum disorders (reviewed in Murphy and Benítez-Burraco 2016). Similarly, some candidates for ASD are important for GABAergic activity, like *MECP2*, involved in DNA methylation, or *GABRB3*, which encodes the β-3 subunit of the GABA receptor A (reviewed in Benítez-Burraco and Murphy 2016). Finally, we wish to highlight genes that are relevant for neural growth and interconnection, such as *DISC1* (related to SZ and encoding a protein involved in neurite outgrowth, cortical development and callosal formation), or *NLGN1* and *SHANK3* (important for establishing functional connections between the cortex and the basal ganglia and found to be mutated in ASD) (see Murphy and Benítez-Burraco 2016 and Benítez-Burraco and Murphy 2016 for details).

Interestingly, one finds common candidates for both disorders which map on to specific aspects of brain function. For instance, an excess of rare novel loss-of-function variants have been found in SZ and ASD in genes important for synaptic function, particularly, in genes coding for neurexin and neuroligin interacting proteins (Kenny et al. 2014). Further, SZ and ASD entail opposite regulation patterns of some specific pathways, like the PI3K signalling pathway, involved in cell survival and brain growth, which is upregulated in some autistics (Philippi et al. 2005, Belmonte and Bourgeron 2006) but is downregulated in schizophrenics (Emamian et al. 2004). Likewise, abnormally high levels of BDNF, a key brain growth factor, have been detected in ASD (Nishimura et al. 2007), whereas SZ usually entails deficiencies in this factor (Palomino et al. 2006). Importantly, some of these common candidates for SZ and ASD are related to abnormal patterns of language development (Wang et al. 2015a), and particularly, to anomalous oscillatory patterns of brain activity. Among them we wish to highlight *ZNF804A* and *PDGFRB*. *ZNF804A* is involved in growth cone function and neurite elongation and plays a role in cortical functioning and neural connectivity (Hinna et al. 2015). Additionally, it coordinates distributed networks belonging to the hippocampus and the prefrontal cortex because of its modulatory role in hippocampal *γ* oscillations (Cousijn et al. 2015). SZ risk polymorphisms of this gene have been related to both phonological and semantic problems (Becker et al. 2012, Nicodemus et al. 2014), whereas ASD risk polymorphisms have been associated with verbal deficits (Anitha et al. 2014). In turn, *PDGFRB* encodes the *β* subunit of the receptor of PDGF, known to be involved in the development of the central nervous system. *Pdgfr-β* knocked-out mice exhibit reduced density of GABAergic neurons in several brain areas, reduced auditory phase-locked *γ* oscillations, alterations of prepulse inhibition, and behavioral disturbances such as reduced social behaviour, impaired spatial memory and problems with conditioning; all of which are aspects are affected in SZ and ASD (Nguyen et al. 2011, Nakamura et al. 2015). Interestingly, *PDGFRA*, which promotes neuronal differentiation, acts downstream of *FOXP2*, the well-known ‘language gene’ (Chiu et al. 2014).

### 4.2. Common candidates for SZ, ASD, and language evolution?

Examining the role played by these types of genes in brain function will directly inform our understanding of the oscillopathic nature of SZ and ASD and, particularly, the nature of the language deficits they entail, as outlined in the previous sections. Ultimately, this approach gives support to the view of SZ and ASD as part of a continuum of cognitive competence encompassing the typically developing faculty of language. Nonetheless, in this paper we have followed a different approach. As noted in the introduction, ample evidence supports the view that modern language was brought about by some reorganizational event in the skull/brain which entailed subtle changes in the wiring, interconnection patterns, and oscillatory behaviour of the hominin brain. As also noted there, modern language is rooted in our species-specific ability to form cross-modular concepts, known to be affected in both SZ and ASD (Lind et al. 2014, Hinzen and Rosselló 2015). In three different, related papers, we have put forth a list of tentative candidates for these changes and for the emergence of our language-readiness (Boeckx and Benítez-Burraco 2014a,b; Benítez-Burraco and Boeckx 2015; see Table 2).

**Table 2.**
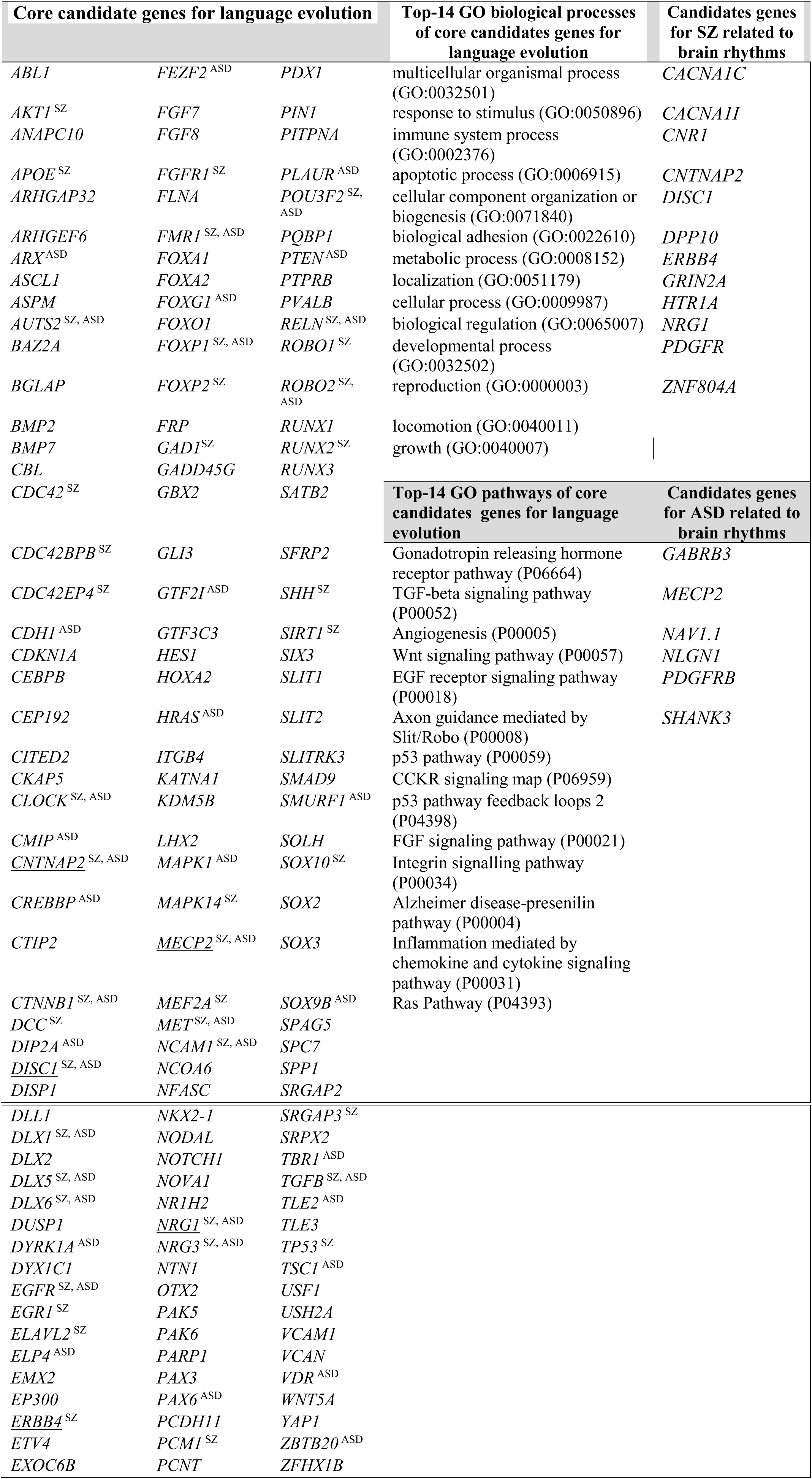
Genes discussed in Section 4. The first column contains core candidates for the evolution of language as posited by Boeckx and Benítez-Burraco (2014a,b) and Benítez-Burraco and Boeckx (2015). Candidates for schizophrenia are marked with ^SZ^, whereas candidates for ASD are marked with ^ASD^. Candidates for language-readiness and for SZ and/or ASD that are thought to be involved in brain rhythmicity have been underlined. Candidates for SZ and/or ASD were identified by literature mining via PubMed (http://www.ncbi.nlm.nih.gov/pubmed). The second column shows a GO classification of the candidates for the evolution of language according to the biological processes (above) or the pathways (below) they contribute to, as provided by Panther (http://pantherdb.org); only the top-25 functions after a Bonferroni correction have been included. The last column comprises genes that have been related to brain rhythms in SZ (above) and ASD (below), as discussed in Benítez-Burraco and Murphy (2016) and Murphy and Benítez-Burraco (2016).

Briefly summarizing the main outcomes of our past research, we have uncovered a set of functionally-related genes involved in aspects of the skull and brain development that bear fixed changes in AMHs in their coding and/or regulatory regions, or that show AMH-specific patterns of expression and/or methylation compared to Neanderthals and/or Denisovans. Specifically, a subset of these genes plays a role in the brain-bone cross-talk important for skull and brain development, like *RUNX2*, some DLX genes (including *DLX1, DLX2, DLX5*, and *DLX6)*, and some *BMP* genes (like *BMP2* and *BMP7)*. Another subset is involved in the regulation of cortical-subcortical interconnection patterns that we believe are important for the externalization of language (speech). This subset encompasses *FOXP2, ROBO1*, and some of the SLITs factors. Finally, some of our candidates (including the ASD-candidate *AUTS2* and some of their partners) provide robust links between the two former subsets of genes. On the whole, these genes map on to specific types of neurons (GABAergic), brain areas (some cortical layers, thalamic nuclei), and processes (importantly, the balance between inhibition and excitation) that are relevant for language processing, supporting the view that modern cognition and language resulted from a set of coordinated changes (Figure 3).

**Figure 3.**
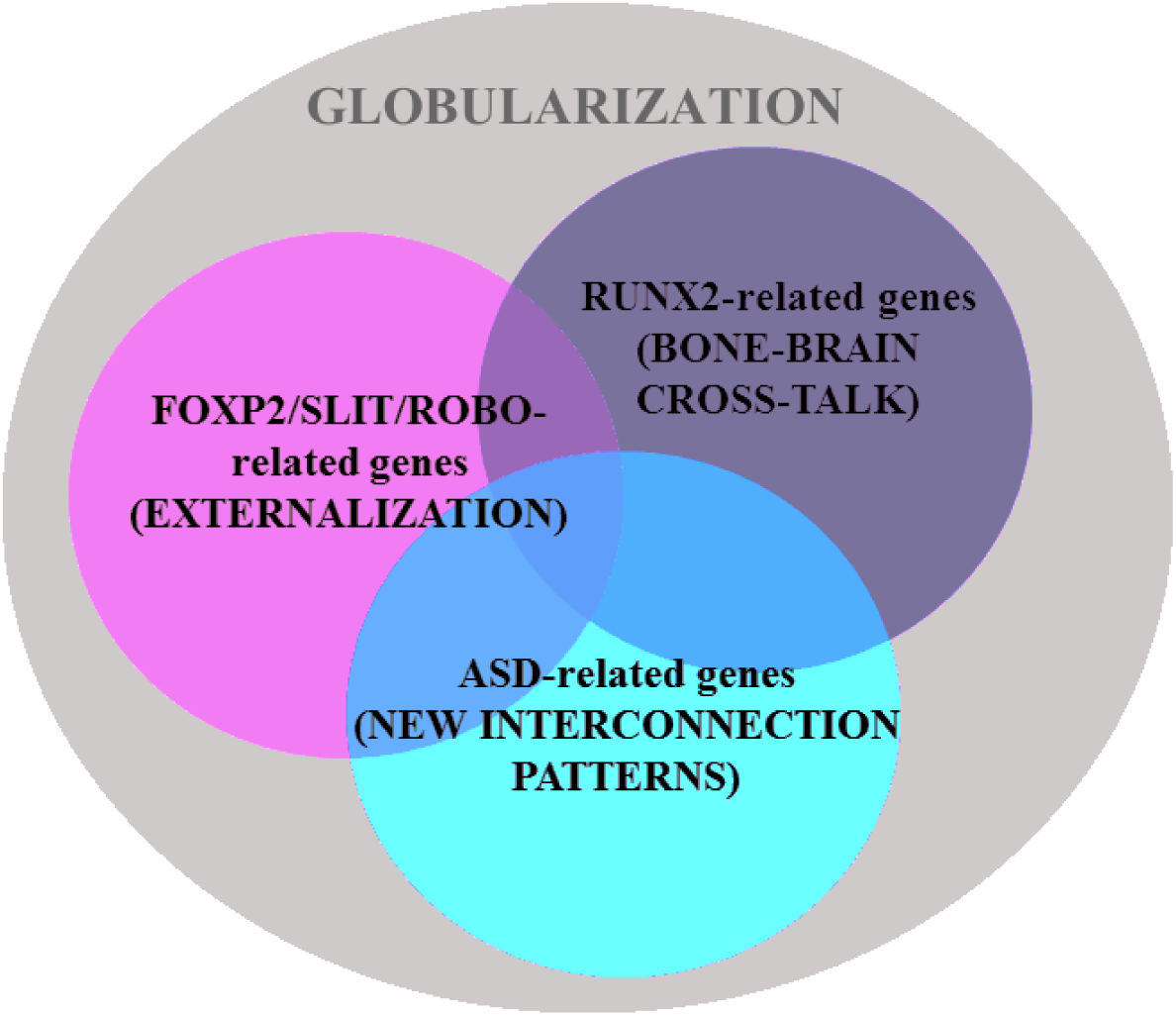
Three sets of functionally-related genes may account for the emergence of language-readiness in our species (based on Boeckx and Benítez-Burraco 2014a,b and Benítez-Burraco and Boeckx 2015).

Interestingly, among the candidates for the globularization of the human skull/brain and the emergence of modern language we have found many candidates for SZ, ASD, or for both conditions (Table 2 and Figure 4). In the second part of this section we will briefly discuss the functions played by these common genes for globularization, SZ and ASD, and will provide some experimental data suggesting that they are dysregulated in these two conditions in areas of the brain important for language processing found and associated with language dysfunction in SZ and ASD.

**Figure 4.**
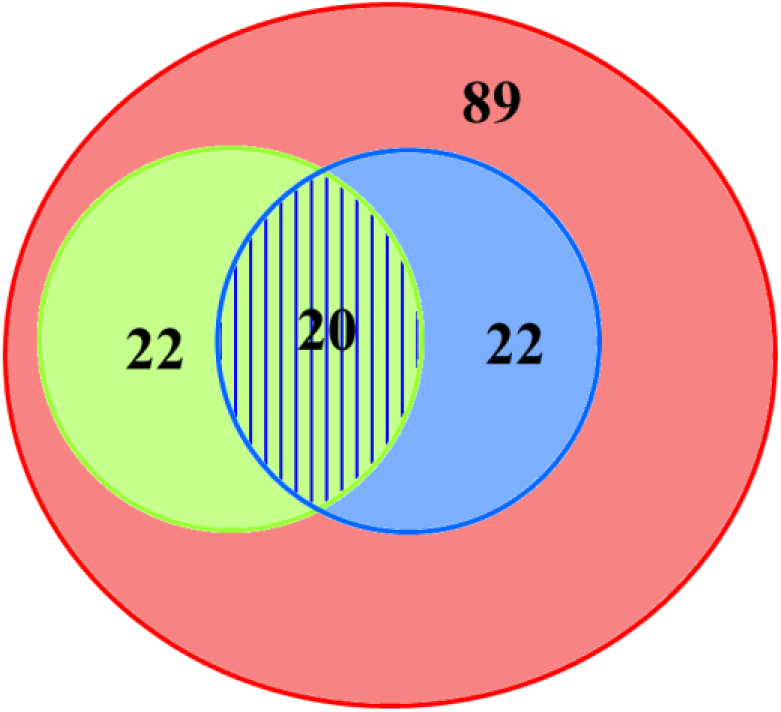
The candidate genes for the emergence of language-readiness in our species (in light red) include many candidates for either SZ (in light green), ASD (in light blue), or both (striped) (see Table 2 for details, and Benítez-Burraco and Murphy 2016 and Murphy and Benítez-Burraco 2016 for discussion).

### 4.3. Common candidates for SZ, ASD and language evolution: a functional characterization

The first one among the candidates for the evolution of language-readiness that are also candidates for both SZ and ASD we will discuss is *CLOCK*, which plays a key role in the regulation of circadian rhythm. Together with other circadian-relevant genes *CLOCK* seems to be involved in the psychopathology of ASD cases entailing sleep disturbances (Yang et al. 2016). It has been hypothesised that changes in the circadian modulation of synaptic function via the synaptic cell adhesion molecules NLGN3, NLGN4 and NRXN1, and the postsynaptic scaffolding protein SHANK3 may decisively contribute to ASD (Bourgeron 2007). Likewise, polymorphisms in *CLOCK* have been associated to SZ in different populations (Zhang et al. 2011, Jung et al. 2014). *CLOCK* interacts with *RUNX2*, one of our core candidates for globularization and a gene showing strong signals of a selective sweep after our split from Neanderthals (Green et al. 2010, Perdomo-Sabotal et al. 2014). *RUNX2* controls different aspects of skull morphology (Stein et al. 2004), but it is also involved in brain development (Jeong et al. 2008, Reale et al. 2013). *RUNX2* is a candidate for SZ (Benes et al. 2007) and interacts with some candidates for ASD, like *SMURF1* (De Rubeis et al. 2014). Additionally, CLOCK is an interactor of several other of our candidates for language-readiness, like *DUSP1* (Doi et al. 2007) and *USF1* (Shimomura et al. 2013). *DUSP1* is a strong candidate for vocal learning, showing a motor-driven expression pattern in the song nuclei of vocal learners (Horita et al. 2010, Horita et al. 2012). Interestingly, DUSP1 interacts with MAPK1 (Choi et al. 2006, Lomonaco et al. 2008), a gene that regulates the transcription of both the FOXP2 target *PLAUR* (Lengyel et al. 1996) and *RUNX2* (Lee et al. 2011), playing as well an important role in osteogenesis in connection with *BMP2* (Ghayor et al. 2009), which is also a member of our core set of genes important for language-readiness. Regarding *USF1*, the regulatory region of this gene shows many fixed or high frequency changes compared to the Denisovan homologue (Meyer et al. 2012). *USF1* regulates synaptic plasticity, neuronal survival and differentiation (Tabuchi et al. 2002, Steiger et al. 2004), and the USF1 protein binds the promoter of *FMR1* (Kumari and Usdin 2001), a strong candidate for Fragile-X syndrome, a condition involving language problems which commonly presents with one or more features of ASD (Kaufmann et al. 2004, Smith et al. 2012). *FMR1* is also a strong candidate for SZ. Accordingly, targets of the FMRP regulon show abnormal protein expression patterns in the brains of schizophrenics (Folsom et al. 2015). Moreover, low levels of FMRP have been associated with lower IQ in people with SZ (Kovács et al. 2013).

As Figure 5 illustrates, String 10 predicts functional interactions between USF1 and another common candidate for SZ, ASD, and language evolution, namely, *CTNNB1*. Like other components of the Wnt/β-catenin pathway, *CTNNB1* is related to both SZ (Levchenko et al. 2015) and ASD (Krumm et al. 2014). This gene encodes β-catenin, a protein involved in cell adhesion. As reviewed in Boeckx and Benítez-Burraco (2014b), CTNNB1 is expected to interact with many of the genes important for globularization and the emergence of language-readiness, like *EP300, CREBB4, SIRT1*, and *CDC42* (predictions based on String 10 data). Interestingly, Ctnnb1 upregulates *Runx2* expression (Han et al. 2014) and the translocation of the Ctnnb1 protein to the nucleus is blocked by an active Slit2/Robo1 signal (Chang et al. 2012). Also interesting is the finding that *CTNNB1* is related to *PCDH11X/Y*, a gene pair that has undergone accelerated evolution in our lineage (Williams et al. 2006) and that has been linked to SZ (Crow 2013), language acquisition delay (Speevak and Farrell 2011), and language evolution due to its role in hemispheric specialisation (Crow 2008, 2013).

**Figure 5.**
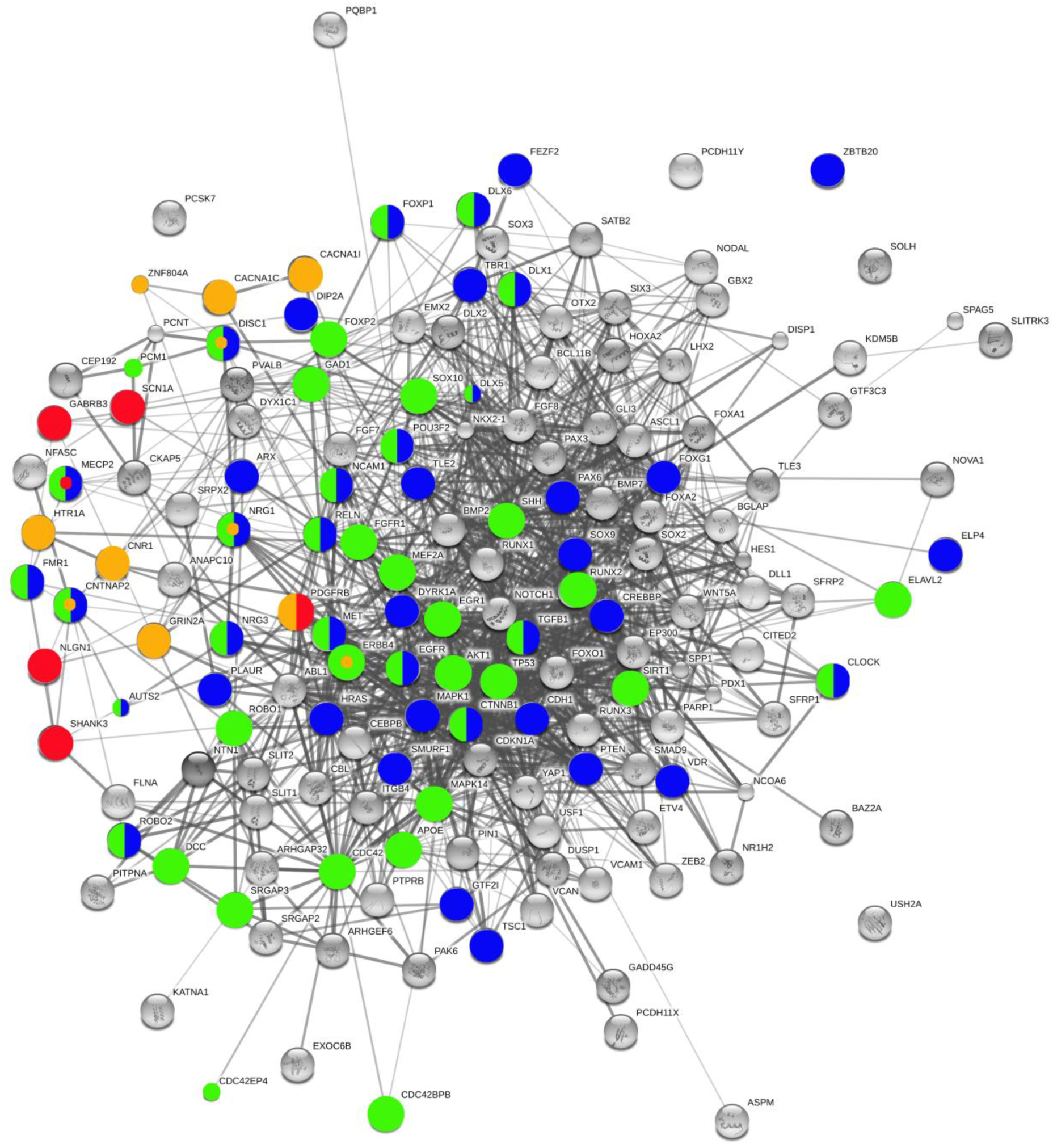
Functional links predicted by String 10 among candidates for the evolution of language and candidate genes for SZ and ASD important for brain rhythmicity. Candidate genes for the evolution of language (as posited by Boeckx and Benítez-Burraco 2014a,b, and Benítez-Burraco and Boeckx 2015) that are also candidates for SZ are colored in light green, whereas those that are also candidates for ASD are colored in blue (otherwise they appear in grey). Genes related to brain rhythms in SZ are colored in orange, whereas those related to brain rhythms in ASD are colored in red. Candidates for language-readiness and SZ and/or ASD that are involved in brain rhythmicity are marked with a small colored dot. The graph displays one candidate for language-readiness SZ and ASD that has not been discussed in the main text: TGFB1. Stronger associations between proteins are represented by thicker lines. The medium confidence value was .0400 (a 40% probability that a predicted link exists between two enzymes in the same metabolic map in the KEGG database: http://www.genome.jp/kegg/pathway.html). String 10 predicts associations between proteins that derive from a limited set of databases: genomic context, high-throughput experiments, conserved co-expression, and the knowledge previously gained from text mining (Szklarczyk et al. 2015). This is why the Figure does not represent a fully connected graph (evidence for additional links are provided in the main text). Importantly, the diagram only represents the potential connectivity between the involved proteins, which has to be mapped onto particular biochemical networks, signaling pathways, cellular properties, aspects of neuronal function, or cell-types of interest that can be confidently related to aspects of language development and function (although see Table 2 and main text for some concerns regarding schizophrenia).

Also of interest is that CLOCK upregulates *VCAM1* (Gao et al. 2014), which encodes a cell surface glycoprotein involved in cell adhesion and the control of neurogenesis (Kokovay et al. 2012). VCAM1 shows a fixed (D414G) change in AMHs compared to Neanderthals/Denisovans (Pääbo 2014, Table S1) and interacts with *NCAM1*, another of our common candidates for SZ, ASD, and language evolution (Vawter et al. 2001, Atz et al. 2007, Zhang et al. 2014). *NCAM1* encodes a cell adhesion protein important for brain development (Fujita et al. 2000, Prag et al. 2002), being involved in axonal and dendritic growth and synaptic plasticity (Hansen et al. 2008, Rønn et al. 2000). Mutations in *Ncam1* impact on working/episodic-like memory (Bisaz et al. 2013). In mice, overexpression of the Ncam1 extracellular proteolytic cleavage fragment impacts on GABAergic innervation and reduces the number of dendritic spines on pyramidal neurons in the prefrontal cortex, affecting long- and short-term potentiation in this area (Brennaman et al. 2011). Finally, we wish highlight that *NCAM1* is also target of both RUNX2 (Kuhlwilm et al. 2013) and FOXP2 (Konopka et al. 2009).

Likewise, three among the four DLX genes highlighted as important for the globularization of the human brain have been related to both SZ and ASD: *DLX1, DLX5*, and *DLX6. DLX1* is differentially expressed across the brain (Johnson et al. 2009) and regulates aspects of the development of the skull and the brain, playing a role in the differentiation and interconnection of thalamic and neocortical neurons (Andrews et al. 2003, Jones and Rubenstein 2004). *Dlx1* downregulation results in reduced glutamatergic input to the hippocampus (Jones et al. 2011), and disturbances in the number of interneuron subtypes and migration patterns in the cortex (Ghanem et al. 2008). *DLX1* is found to be downregulated in ASD (Voienagu et al. 2011, McKinsey et al. 2013) and in the thalamus of schizophrenics (Kromkamp et al. 2003). Among the DLX1 interactors, one finds another of our candidates, *ROBO2*, involved in thalamocortical axons development, which represent the major input to the neocortex and which modulate cognitive functions (López-Bendito et al. 2007, Marcos-Mondéjar et al. 2012). Also *ROBO2* is a candidate for ASD (Suda et al. 2011) and SZ (Potkin et al. 2009, 2010), although it has been associated as well to dyslexia (Fisher et al. 2002), speech-sound disorder and reading (Stein et al. 2004), and expressive vocabulary growth in typically developing individuals (St Pourcain et al. 2014). *ROBO2* shows a specialized expression pattern of in some of the areas involved in song learning and performance in songbirds (Wang 2011). Concerning *DLX5* and *DLX6*, they encode bone morphogenetic factors important for skull and brain development (Kraus and Lufkin 2006, Wang et al. 2010). As was true of *DLX1*, they regulate the migration and differentiation of precursor cells that give rise to GABA-expressing neurons in the forebrain (Cobos et al. 2006) and play a role in thalamic development too (Jones and Rubenstein 2004). Both *DLX5* and *DLX6* are believed to contribute to the aetiopathogenesis of ASD (Nakashima et al. 2010). Likewise Dlx5/6(+/-) mice exhibit abnormal patterns of *γ* rhythms resulting from alterations in GABAergic interneurons, particularly in fast-spiking interneurons, which are a hallmark of SZ (Cho et al. 2015). Interestingly, an ultraconserved cis-regulatory element affects to both *DLX5* and *DLX6* and contributes to modulate their role in regulating the migration and differentiation of GABA-expressing neurons in the forebrain: This regulatory element has been found mutated in people with ASD (Poitras et al. 2010) and it is bound by GTF2I, encoded by one of the genes commonly deleted in Williams-Beuren syndrome and a candidate for ASD too (Malenfant et al. 2012). *DLX5* is reported to be modulated by MECP2 (Miyano et al. 2008). *MECP2* is the main candidate for Rett syndrome, a neurodegenerative condition entailing language loss, problems for motor coordination, and autistic behaviour (Uchino et al. 2001, Veenstra-VanderWeele and Cook 2004). *MECP2* is critical for normal function of GABA-releasing neurons (Chao et al. 2010). The gene is also found to be mutated in SZ (Cohen et al. 2002, McCarthy et al. 2014). On the whole, these evidence give additional support to the view that dysfunction in GABA signalling mediates ASD-like stereotypies and Rett syndrome phenotypes (Chao et al. 2010), but also SZ (Fazzari et al. 2010). In addition, DLX5 interacts with two of our key candidates for language evolution. Hence, it regulates the expression of *RUNX2* (Jang et al. 2011), whereas the expression of both *Dlx5* and *Dlx6* is regulated by Foxp2 via *Shhrs*, a noncoding RNA (Vernes et al. 2011). Not surprisingly, *Foxp2* and *Dlx5* are expressed in the same parts of the amygdala in rats and nonhuman primates, and in almost the same neuronal populations of the striatum (Kaoru et al. 2010), a subcortical component that plays a key role in language processing which has changed during recent human evolution (Raghanti et al. 2015).

It is not surprising, then, that we have found that several interactors of FOXP2 that are important for globularization and language evolution are also candidates for both SZ and ASD. Among them we wish to highlight *FOXP1*, *POU3F2*, *MET*, *CNTNAP2*, and *DISC1*. The protein encoded by *FOXP1* binds to the protein FOXP2 and form a functional heterodimer (Li et al. 2004), playing a role in postmigratory neuronal differentiation (Ferland et al. 2003). The rat *Foxp1* shows altered expression in animals exhibiting a hypofunction of the NMDA-type glutamate receptor (known to exacerbate SZ symptoms in patients) (Ingason et al. 2015). Mutations in the gene have been related to ASD, but also to other cognitive disorders, including language impairment and intellectual disability (Hamdan et al. 2010, Horn et al. 2010). Regarding POU3F2, this protein binds a specific site within intron 8 of *FOXP2* (Maricic et al. 2013). Interestingly, Neanderthals and Denisovans bore an ancestral allele of the binding site which was more efficient in activating transcription (Maricic et al. 2013). *POU3F2* is thought to be involved in dopamine and serotonin synthesis in several brain areas (Nasu et al. 2014) and to regulate as well the migration and the identity of neocortical upper-layer neurons (McEvilly et al. 2002, Sugitani et al. 2002). Sequence and copy number variations affecting *POU3F2* have been found in people with SZ and ASD, but also in patients suffering from developmental and language delays (Huang et al. 2005, Potkin et al. 2009, Lin et al. 2011). With regards to *MET*, *CNTNAP2* and *DISC1*, these are well-known targets of FOXP2 (Vernes et al. 2008, Mukamel et al. 2011, Walker et al. 2012). *MET* encodes a membrane receptor tyrosine kinase involved in axon terminal outgrowth and synaptogenesis, dendritic branching, spine maturation, and excitatory connectivity (Judson et al. 2011). The gene has been found important for neocortical and cerebellar growth and maturation (Campbell et al., 2006), and particularly, for the regulation of the glutamatergic synapse maturation and cortical circuit function (Peng et al. 2016). *MET* influences SZ risk and neurocognition (Burdick et al. 2010) and it is a robust candidate for ASD, because dysregulated MET signaling has been hypothesised to alter cortical thickness and forebrain maturation and connectivity in this condition (Hedrick et al. 2012, Rudie et al. 2012, Peng et al. 2016), resulting in abnormal patterns of fMRI activation and deactivation in response to social stimuli (Rudie et al. 2012). As noted above, *CNTNAP2* encodes a protein associated with K+ voltage-gated channels found in pyramidal cells that are mostly innervated by GABAergic interneurons (Inda et al. 2006). *CNTNAP2* regulates axonogenesis in conjunction with ROBO and SLIT factors (Banerjee et al. 2010), and it is also important for synaptogenesis (Dean et al. 2003), and it plays a role as well in regulating brain connectivity and cerebral morphology (Scott-Van Zeeland et al. 2010, Tan et al. 2010, Dennis et al. 2011) and dendritic arborization and spine development (Anderson et al. 2012). Additionally, it seems to contribute to vocal behaviour: *Cntnap2* is highly expressed in the males of the zebrafinch in the lateral magnocellular nucleus of the anterior nidopallium, but also in the robust nucleus of the arcopallium (Panaitof *et al.*, 2010). The gene is a candidate for ASD (Alarcón et al. 2008, Bakkaloglu et al. 2008) and language impairment in SZ (Poot 2015), although it has been related as well to intellectual disability (Gregor et al. 2011), and different types of language disorders, including SLI (Newbury et al. 2011), dyslexia (Peter et al. 2011), child apraxia of speech (Worthey et al. 2013), and variants of language delay and language impairment (Petrin et al. 2010, Sehested et al. 2010). Non-pathogenic polymorphisms of *CNTNAP2* have been proved to affect language development in the normal population (Whitehouse et al. 2011, Whalley et al. 2011, Kos et al. 2011). Interestingly, the AMH protein bears a fixed change (Ile345Val) compared to the Denisovan protein (Meyer et al. 2012). Moreover, in mice *Cntnap2* is found among Auts2 regulatory targets (Oksenberg et al. 2014). *AUTS2* is the gene that displays the strongest signal of a selective sweep in AMHs compared to Neanderthals (Green et al. 2010, Oksenberg et al. 2013) and a robust candidate for ASD (Oksenberg and Ahituv 2013) and schizoaffective disorder (Hamshere et al., 2009), but also for bipolar disorder (Hattori et al. 2009), differential processing speed (Luciano et al. 2011), dyslexia (Girirajan et al. 2011), and intellectual disability and developmental delay (Oksenberg and Ahituv 2013). Mutations in *AUTS2* also impact skull development (Talkowski et al. 2012, Beunders et al. 2013, Oksenberg et al. 2013). Finally, concerning *DISC1*, this is one of the more robust candidates for SZ and other cognitive and brain disorders like ASD (Johnstone et al. 2011, Bradshaw and Porteous 2012, Narayan et al. 2013, Kenny et al. 2014). *DISC1* is known to play a role in neurite outgrowth, cortical development and callosal formation (Brandon and Sawa 2011, Osbun et al. 2011), and particularly in regulating microcircuit architecture in the dentate gyrus through its control of neurogenesis (Reif et al. 2007, Kim et al. 2012). Mutations in *Disc1* causes a deficit in spatial working memory (Kvajo et al. 2008). For our purposes it is useful to note that pathological disruptions of information flow through the neural microcircuits responsible for adaptive behaviors have been related to changes in synaptic plasticity, either to short-term synaptic plasticity alterations (in the case of SZ) or to a combination of both short-term and long-term synaptic plasticity alterations (in the case of ASD) (Crabtree and Gogos 2014). Moreover, transgenic mice expressing a truncated version of *Disc1* show an altered pattern of *θ* burst-induced long-term potentiation (and, ultimately, of long-term synaptic plasticity) (Booth et al. 2014). During the formation of excitatory-inhibitory synapses by cortical interneurons DISC1 plays an inhibitory effect on NRG1-induced ERBB4 activation and signalling (Seshadri et al. 2015). One interactor of DISC1 is *PCNT*, a candidate for dyslexia (Miyoshi et al. 2004). *PCNT* is mentioned by Green et al. (2010) as being among the top ten genes showing non-synonymous and non-fixed substitution changes in their coding sequences compared to Neanderthals.

As noted in Table 2, NRG1 (discussed above) and NRG3 are commonly cited as candidates for both SZ and ASD (Kao et al. 2010, Hatzimanolis et al. 2013, Hou et al. 2014, Tost et al. 2014, Shen et al. 2015, Yoo et al. 2015, Zeledón et al. 2015). Together with its receptor ERBB4 (also discussed above), NRG1 is involved in the enhancement of synchronized oscillations in the prefrontal cortex, known to be reduced in SZ (Fisahn et al. 2009, Hou et al. 2014). Risk polymorphisms of *NRG1* are associated with language performance in people with bipolar disorder (Rolstad et al. 2015) and correlate with reduced volumes of the left superior temporal gyrus (a robust imaging finding in SZ) (Tosato et al. 2012), a region involved in language processing (Aeby et al. 2013). Concerning *NRG3*_*1*_ it is known to influence prefrontal cortex activation during working memory performance (Tost et al. 2014). Like *NRG1*, *NRG3* is also a candidate for atypical neurodevelopmental outcomes (Blair et al. 2015) and speech delay (van Bon et al. 2011). And in conjunction with *NRG1*, *NRG3* contributes to regulate the migration of GABAergic interneurons from ganglionic eminences to their cortical targets (Li et al. 2012).

Finally, we wish to highlight *RELN*, involved in neuronal migration and also a common candidate for both SZ and ASD (Ashley-Koch et al. 2007, Holt et al. 2010, Wang et al. 2014, Li et al. 2015); and *EGFR*, which encodes a receptor for members of the epidermal growth factor family and which is also one of RUNX2’s targets (Kuhlwilm et al. 2013) and a candidate for SZ (Benzel et al. 2007) and ASD (Russo, 2014, 2015). *EGFR* is functionally linked to *VCAN*, a gene which encodes a protein involved in neuronal attachment, neurite outgrowth, and synaptic transmission (Xiang et al. 2006), which shows a fixed (N3042D) change in AMHs compared to Neanderthals and Denisovans (Pääbo, 2014; Table S1).

### 4.4. Common candidates for SZ, ASD, and language evolution: a global view

We expect that the genes we highlight above are interconnected at some functional level and map on to specific pathways, signaling cascades, or aspects of brain development and function of interest for language processing and the aetiopathology of SZ and ASD. In silico analyses offer promising insights. Accordingly, String 10 (www.string-db.org) predicts quite robust links between most of these genes (Figure 6). Likewise, ontology analyses by Panther (www.pantherdb.org) suggests that some of them are part of the same signaling pathways. Hence, *EGFR*, *NRG1*, and *NRG3* are part of the EGF receptor signaling pathway (P00018), whereas *EGFR* and *CTNNB1* are part of the cadherin signaling pathway (P00012). The available literature offers some additional interesting findings. Accordingly, and as noted above, some of these genes are involved in the development of GABAergic circuitry. Specifically, in the developing ventral forebrain of mice the same ncRNA recruits transcription of*Mecp2, Dlx5*, and *Dlx6* for properly formatting GABAergic circuits in the hippocampus and the dentate gyrus (Bond et al. 2009). Likewise, *Dlx1* and *Robo2* (but also some other candidates for globularization that have been associated to either SZ or ASD, like Erbb4), are involved in the regulation of tangential migration to specific brain areas, like the olfactory bulb (Long et al. 2007). Finally, *DISC1*, *NRG1*, *CTNNB1*, and *RELN* (and also the PI3K/AKT signaling pathway discussed below) are required for neurogenesis and migration of neurons as part of the cell non-autonomous neurodevelopmental mechanisms (see Figure 7 and Narayan et al. 2013 for details).

**Figure 6.**
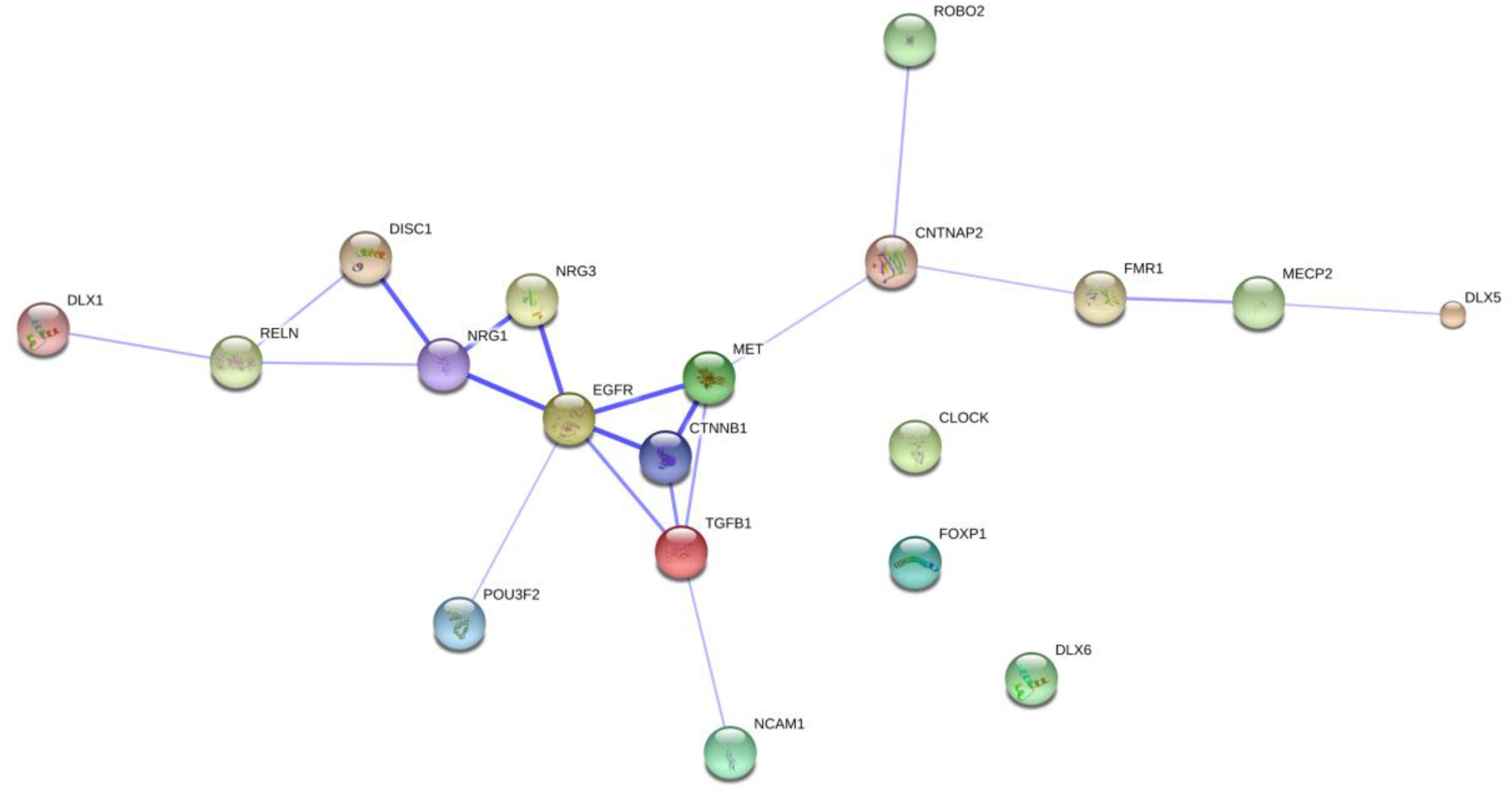
Functional links predicted by String 10 among the genes that are candidates for the evolution of language, SZ and ASD as discussed in the text. The caveats noted for Figure 5 crep apply.

**Figure 7.**
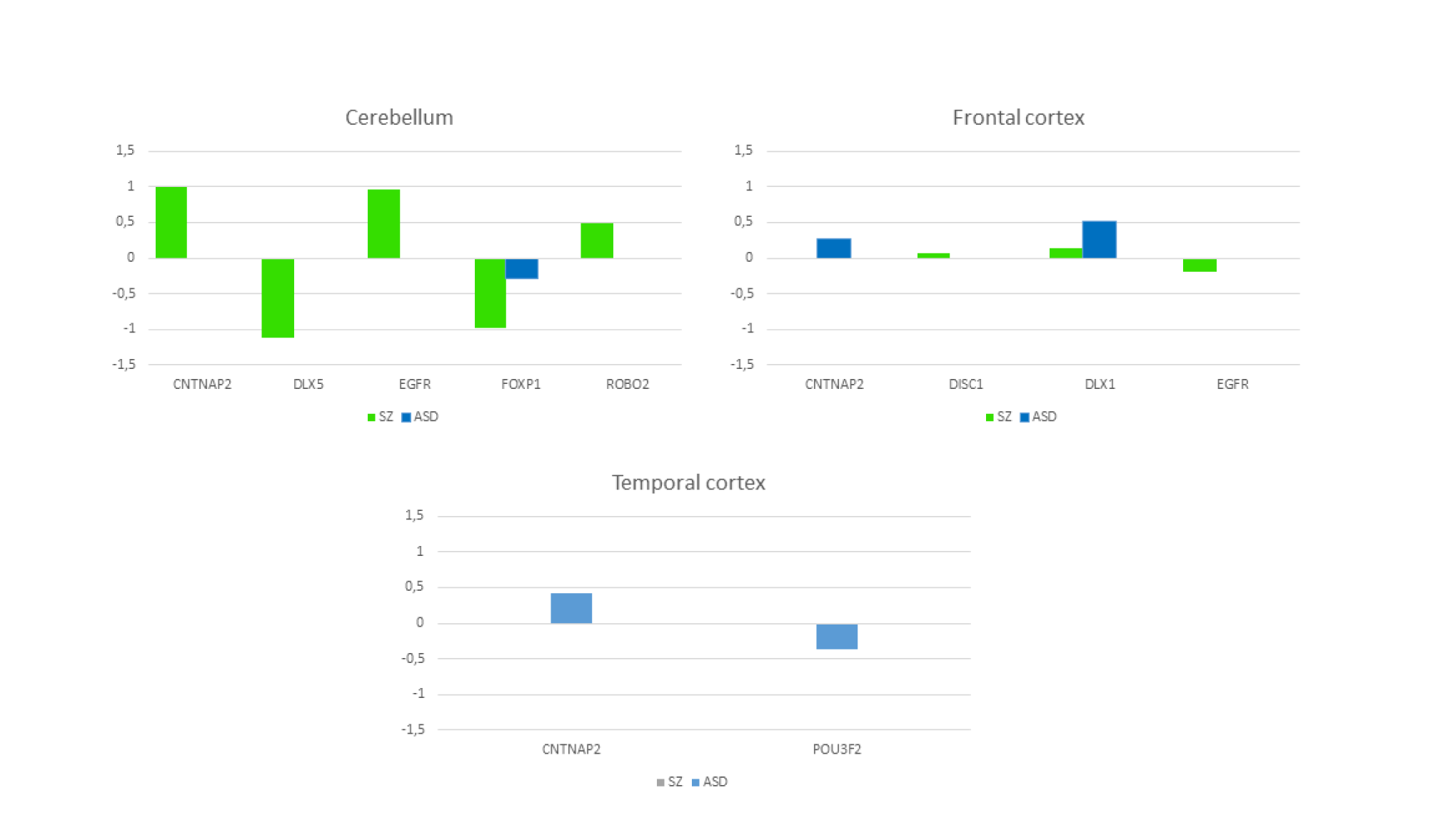
Comparative expression profiles of common candidates for SZ, ASD, and language-readiness in the human brain. The data were obtained from microarray expression datasets available on the Gene Expression Omnibus database (GEO datasets, http://www.ncbi.nlm.nih.gov/gds): GSE28521 (Voineagu et al. 2011) for ASD patients, and GSE4036 (Perrone-Bizzozero unpublished results), GSE17612 (Maycox et al. 2009), and GSE21935 (Barnes et al. 2011) for SZ patients (for the cerebellum, frontal cortex, and superior temporal cortex, respectively). Datasets were analysed using GEO2R (available at http://www.ncbi.nlm.nih.gov/geo/geo2r), comparing the original submitter-supplied data in a pairwise fashion (i.e. patients-versus-controls) for each selected brain area. Details on algorithm dataflow and technical specifications are available in Barrett et al. (2013). In our analysis, the p-value was adjusted using Bonferroni-Hochberg correction for false discovery rate (FDR). Data are shown as log transformation of fold changes (logFC) between patients (ASD or SZ) and corresponding controls. Only genes with probes of p ≤ 0.05 are considered.

As noted in section 1 and 2 above, Crespi and Badcock (2008) have recently put forth the hypothesis that SZ and ASD share a common aetiology. In particular, after reviewing ample evidence showing that these two conditions exhibit diametric phenotypes – from cognitive (dys)abilities, to brain anatomy and function, to neurodevelopmental paths – they hypothesized that these differences may result from opposite alterations of genomic imprinting. Hence, SZ would involve increased relative bias towards effects of maternally expressed genes, whereas ASD would involve biases towards increased relative effects from imprinted genes with paternal expression. We have searched our candidates in Gene Imprint http://www.geneimprint.com/) and found that only *DLX5* is maternally imprinted in humans. Nonetheless, we hypothesize that common candidates for SZ and ASD show dysregulated (and perhaps opposite) patterns of expression in the brain. There are not many studies examining this possibility, but recent large-scale studies aimed to detect opposing risks of SZ and ASD depending on normal variation provide support to the hypothesis that diametric gene-dosage effects may contribute to these two disorders; for example, those examining adjusted body size at birth, a proxy for diametric gene-dosage variation in utero (Byars et al. 2014). Consequently, we searched for the expression profiles of our common candidates for SZ, ASD, and globularization in human brain tissues, and found that the genes for which we could find confident expression values are dysregulated in the cerebellum, but also in the frontal cortex and the temporal cortex. Nonetheless, because of the scarcity of the data, we could not check whether they also show opposite expression patterns in these areas (Figure 7).

## 5 Conclusions

Next generation sequencing technologies have exponentially increased the number of genes related to cognitive diseases like SZ and ASD. In truth, the polygenism seen in both conditions is somewhat commensurable with the polygenism displayed in language more generally, which we also need to understand if we want to gain an accurate view of how language grows and develops in the child. At the same time, these technologies have provided us with a detailed view of the changes in the human genome that were brought about during and after our split from extinct hominins and that seemingly contributed to the emergence of our species-specific cognitive abilities, language being the most outstanding. Bridging this gap between gene change, the emergence of new brain functions, and the achievement of enhanced cognitive abilities is a real challenge. As noted in the introduction, the gap between gene mutations, neuropathology, and cognitive dysfunction is equally broad. Interestingly, SZ and ASD exhibit a notable and stable prevalence across cultures and epochs, whereas they are absent or tenuous in other close species, like great apes. Overall, this is suggestive of a deep, robust link between cognitive impairment and cognitive evolution, and specifically, between language disorders and the emergence of our language-readiness.

0ur main concerns have been to use SZ and ASD to construct a robust model of linguistic computation in the brain, to construct successful oscillatory endophenotypes of these conditions, and to try and bridge the gap between language dysfunction and the genome. In working towards the latter goal, we have focused on the gene changes that are believed to be important for the evolution of language, but also in how language is processed in the impaired brain, with a focus on brain oscillations due to their putative role in cognitive functioning (see Figure 8). In truth, both aspects are interrelated, since some genes that are important for brain rhythmicity are candidates for SZ and/or ASD, but they are also related to the changes that modified brain wiring during human speciation (see Benítez-Burraco and Murphy 2016 and Murphy and Benítez-Burraco 2016 for details).

**Figure 8.**
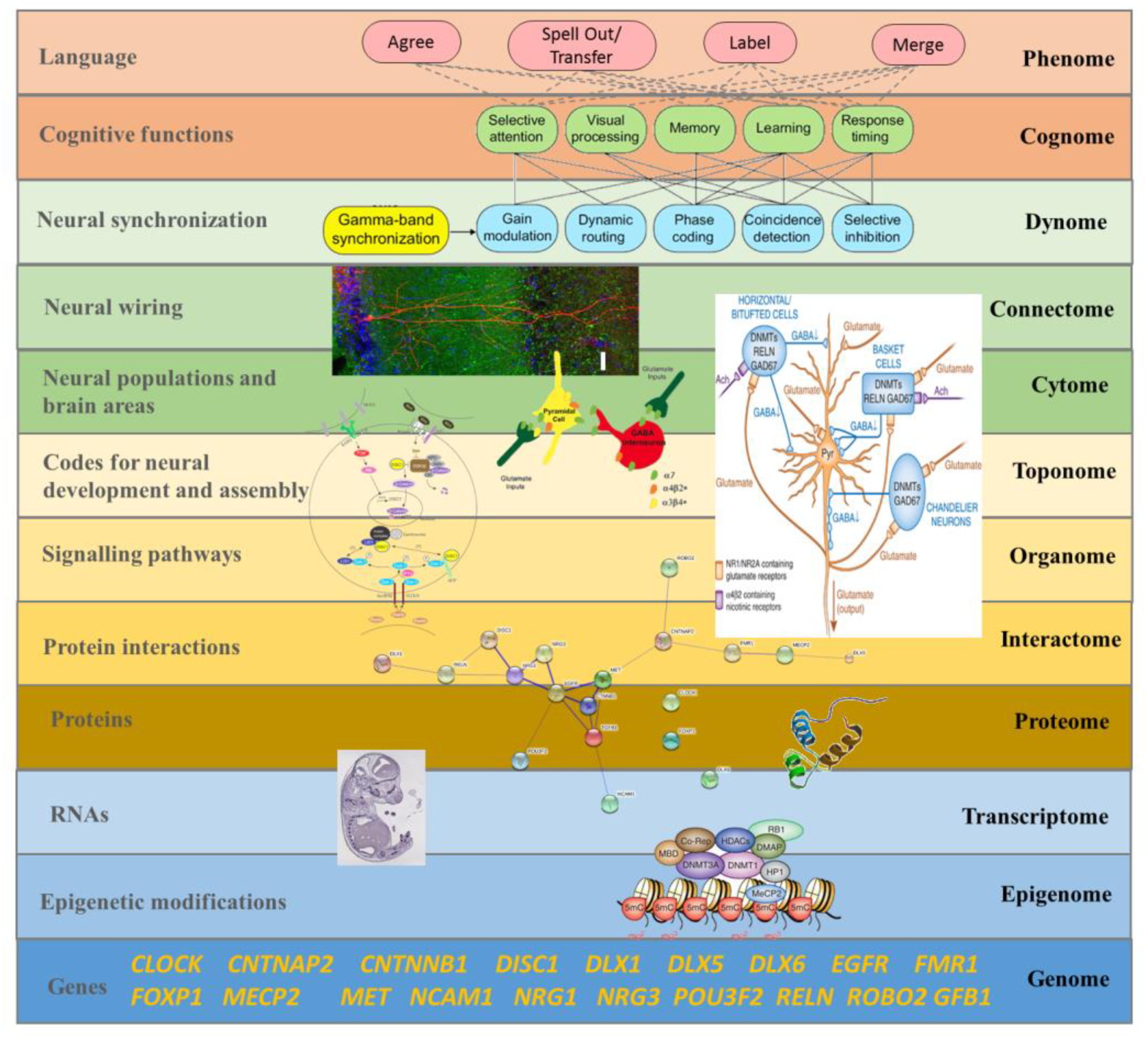
A multilevel approach to language deficits in SZ and ASD from an oscillatory perspective. Understanding language problems in SZ and ASD demands a systems biology approach that seeks to unravel the nature and the links between all the biological factors involved. The figure shows one possible line of research focused on brain oscillations, although many others will need to be explored in the future. As noted in the main text, there exist several common candidates for SZ and ASD (at the bottom of the Figure) that are related to the skull and brain changes that occurred during recent human evolution. As the String 10 network displayed directly above them shows, most of these genes are functionally interrelated and belong to key signalling pathways and regulatory networks important for brain development and function. The network in the middle of the figure shows the central role played by one of these genes (DISC1) in several pathways relevant for neuronal migration and positioning: the Reelin cascade, the NRG1/ErbB signaling pathway, and the PI3K/Akt signalling, known to be disturbed in SZ and ASD. As also noted in the text, disturbances in GABAergic mediator systems may contribute to the altered γ activity observed in both conditions, ultimately impacting on language processing (on the top of the Figure), although the exact role played by these basic cognitive operations in language processing is still unknown (dashed lines). This is a composite Figure elaborated by the authors. The oscillomic-cognomic aspects of γ-oscillations (at the top of the figure) are from Bosman et al. 2014. The micrography of the single hippocampal CA1 pyramidal neuron is from http://basulab.us/research/goals. The schematic view of the hippocampal GABAergic neurons (below) is from Feduccia et al. (2012). The scheme of the role of DISC1 in cell non-autonomous mechanisms regulating cortical development is from Narayan et al. (2013). The String 10 network is from Figure 5 (this paper). The schematic view of the structure of the DLX1 protein has been taken from www.uscnk.com. The in situ hybridization of Dlx1 in E10.5 mouse embryo is from Panganiban and Rubenstein (2002). The schematic representation of the transcriptionally inactive promoter is from Grayson et al. (2013). Finally, the large scheme on the right shows the effect of DNA methyltransferase overexpression on GABAergic neurons and also comes from Grayson et al. (2013); of interest is that GABAergic promoter downregulation is observed in SZ, resulting in increased levels of some DNA methyltransferases (like DNMT1 and 3A), and reduced GAD67, RELN and a variety of interneuron markers.

As we hope to have shown, this connection between language evolution and language impairment is a robust one – more robust perhaps than previously believed. Candidates for either SZ or ASD, or for both diseases, are overrepresented among the genes that seem to have played a central role in the brain changes that prompted our ability to learn and use languages, implemented via the oscillomic models in Figures 1 and 2. As noted, they map on to specific neuronal types (e.g. GABAergic), particular brain areas (e.g. frontal cortex), particular physiological processes (the balance between inhibition and excitation), specific developmental processes (inter and interhemispheric axon pathfinding, or brain lateralization), and particular cognitive abilities (cross-modular thought), all of which are aspects known to be impaired in SZ and ASD, but are also aspects known to have changed during human evolution. Also pertinent in this respect is the observation that genes important for language evolution that are known to be involved in the aetiopathogenesis of both SZ and ASD are dysregulated in brain areas known to be involved in language processing, giving support to the view that SZ and ASD are cognitive diseases resulting for subtle changes in the same aetiological factors. On the whole, we feel justified in hypothesising that the same genes that cause SZ and ASD are involved in aspects of brain development and function that are important for language, but also that the aspects of brain development and function that changed during our recent evolution are quite sensitive to damage in modern populations. Importantly, this damage results in cognitive impairment, although we wish stress that this applies to specific kinds of cognitive impairment only. As noted in the introduction, recently evolved neural networks are more sensitive to damage because they lack robust compensatory mechanisms (Toro et al. 2010), and so we would predict that these mechanisms underlie the hypothesized impaired rhythms (for instance, low to middle γ). It seems, then, that the transition from an ape-like cognition to a human-specific cognition (entailing brain wiring modifications and the emergence of new patterns of sub- and cross-cortical neural oscillations) made the latter substantially vulnerable to damage, plausibly because this process (involving gene mutations, but also cultural changes and demographic bottlenecks) uncovered cryptic variation and de-canalized primate cognition, which is particularly robust after millions of years of stabilizing selection (see Gibson 2009 for details). If we are on the right track, this may explain the high prevalence of SZ and ASD among modern populations, along with their overlapping aetiologies.

We wish to end by emphasising that our results should encourage any research program aimed at translating linguistic computation into a specific code of brain activity patterns, like the kind proposed in Murphy (2015a) and discussed here. Likewise, it is important to qualify the common-lay view that cognitive disease is the ‘price we paid for language’ (e.g. Crow 1997 on SZ). More specifically, conditions like SZ and ASD are the consequences of having gained a more globally efficient ability for recursive oscillatory embedding. Seeing language deficits in cognitive disease as specific alterations of the cognome-dynome cross-talk seems to us a promising direction for future research, particularly if other levels of description are also considered (Figure 8). Furthermore, connectomopathies like SZ and ASD should be better construed as restricted areas within the adaptive landscape of the human cognitive phenotype, resulting from the abnormal interaction of the factors that regulate the development of the human brain and cognition (see Benítez-Burraco 2016 for details). Finally, we further expect that the present proposal will help achieve more robust endophenotypes of SZ and ASD (and of language deficits in both conditions), in turn aiding clinical linguists and psychiatrists in improving their diagnosis and therapeutic approaches to these disorders.

## Acknowledgements

The authors wish to sincerely thank Wanda Lattanzi (Università Cattolica del Sacro Cuore, Rome, Italy) for providing the curated data of the GEO datasets employed in Figure 7. Preparation of this work was supported in part by funds from the Spanish Ministry of Economy and Competitiveness (grant numbers FFI2014-61888-EXP and FFI-2013-43823-P to ABB) and in part by an Economic and Social Research Council scholarship (grant number 1474910 to EM).

